# Bidirectional, unlike unidirectional transport, allows transporting axonal cargos against their concentration gradient

**DOI:** 10.1101/2021.01.27.428531

**Authors:** Ivan A. Kuznetsov, Andrey V. Kuznetsov

## Abstract

Even though most axonal cargos are synthesized in the soma, the concentration of many of these cargos is larger at the presynaptic terminal than in the soma. This requires transport of these cargos from the soma to the presynaptic terminal or other active sites in the axon. Axons utilize both bidirectional (for example, slow axonal transport) and unidirectional (for example, fast anterograde axonal transport) modes of cargo transport. Bidirectional transport seems to be less efficient because it requires more time and takes more energy to deliver cargos. In this paper, we studied a family of models which differ by the modes of axonal cargo transport (such as anterograde and retrograde motor-driven transport and passive diffusion) as well as by the presence or absence of pausing states. The models are studied to investigate their ability to describe axonal transport against the cargo concentration gradient. We argue that bidirectional axonal transport is described by a higher-order mathematical model, which allows imposing cargo concentration not only at the axon hillock but also at the axon terminal. The unidirectional transport model allows only for the imposition of cargo concentration at the axon hillock. Due to the great lengths of the axons, anterograde transport mostly relies on molecular motors, such as kinesins, to deliver cargos synthesized in the soma to the terminal and other active sites in the axon. Retrograde transport can be also motor-driven, in which case cargos are transported by dynein motors. If cargo concentration at the axon tip is higher than at the axon hillock, retrograde transport can also occur by cargo diffusion. However, because many axonal cargos are large or they assemble in multiprotein complexes for axonal transport, the diffusivity of such cargos is very small. We investigated the case of a small cargo diffusivity using a perturbation technique and found that for this case the effect of diffusion is limited to a very thin diffusion boundary layer near the axon tip. If cargo diffusivity is decreased in the model, we show that without motor-driven retrograde transport the model is unable to describe a high cargo concentration at the axon tip. To the best of our knowledge, our paper presents the first explanation for the utilization of seemingly inefficient bidirectional transport in neurons.

## 1. Introduction

To ensure reliable operation over decades, neurons need to constantly replace their functional components. In neurons, the delivery of new components to distal parts of an axon is accomplished by active transport that requires the involvement of molecular motors. It is likely that most neuronal proteins are replaced once every several weeks by new proteins arriving from the soma (Misgeld and Schwarz 2017; Goldberg 2003). The question of how axonal cargos are delivered to various axonal sites has been extensively investigated (Miles and Keener 2017; Bressloff and Karamched 2016; Lee and Mitchell 2015; Karamched and Bressloff 2017; Newby and Bressloff 2010; Miles et al. 2018).

Cytosolic proteins are transported from the soma, where they are synthesized, to active sites, e.g., the presynaptic terminal, by slow component-b (SCb). SCb is involved in the transport of ~200 proteins (Roy et al. 2008). The average rate of SCb-transport is ~2-8 mm/day (0.023-0.093 μm/s). One of the proteins transported in SCb is α-synuclein (α-syn), which is mostly known for its involvement in Parkinson’s disease (PD) (Charvin et al. 2018; Shahmoradian et al. 2019). In a healthy neuron α-syn predominantly exists as a monomer (Lashuel et al. 2013), and hereafter when we write α-syn we mean α-syn monomer unless we specify otherwise. α-syn is synthesized in the soma and is then transported to the synapse, mostly in SCb (Jensen et al. 1998; Jensen et al. 1999; Tang et al. 2012). In healthy neurons α-syn predominantly localizes in the presynaptic terminal (Li, J. Y. et al. 2002; Fortin et al. 2005; Burre 2015). As for any SCb protein, α-syn motion is characterized by periods of fast transport, pauses, and changes in movement direction (Yang et al. 2010). Fast anterograde and retrograde motions of α-syn are propelled by kinesin and dynein motors, respectively (Utton et al. 2005).

An intriguing question is why cytosolic proteins, such as α-syn (Roy 2016), are moved in SCb. SCb includes both anterograde and retrograde components, although the net flux of these proteins is anterograde. Transporting cargos using molecular motors requires energy (Maday et al. 2014). Bidirectional transport takes more energy to move cargo by the same distance compared to unidirectional transport. Additionally, bidirectional transport (especially slow axonal transport) is slower than unidirectional transport. Slow axonal transport along human motor neurons can take up to a year. This may negatively affect neurons’ ability to supply presynaptic terminals with key proteins, as slow transport acts as a bottleneck and may predispose them to proteolytic degradation in transport. Indeed, the cellular environment is quite hostile to most proteins, and most of them have short life, days or weeks at the most (Goldberg 2003). For some unknown reason, neurons use slow axonal transport, which is inherently bidirectional, to deliver three times more protein to the presynaptic terminal than by fast axonal transport (Maday et al. 2014). Why has evolution chosen the mode of transport that includes a retrograde component?

Back-and-forth movements of cargos with overall net directionality toward a certain destination seemingly constitute an inefficient transport mechanism. Possible explanations for the utilization of bidirectional transport include facilitating the maneuvering of molecular motors around obstacles on microtubules (MTs) and providing an error correction mechanism for the delivery of cargo to a proper location (Hancock 2014). It was also suggested that the purpose of bidirectional transport (especially of slow axonal transport) is the dynamic recruitment and redistribution of cargos in response to changing metabolic needs (Brown 2016).

Unlike previously suggested hypotheses, our hypothesis is that bidirectional transport in neurons allows for the movement of cargos from a location with a lower cargo concentration to a location with a higher cargo concentration. We refer to this situation as cargo transport against its concentration gradient; it requires cargo transport against its diffusion-driven flux. From a physics perspective, transport against a concentration gradient is not unusual; it requires energy to allow the corresponding decrease in entropy. Ion pumps, for example, transport ions through pores against a gradient using energy supplied by ATP hydrolysis. Similarly, ATP hydrolysis drives the stepping of the molecular motors and the generation of force allowing transport against cargo concentration. Situations where cargos must be transported against their concentration gradient are common in neurons. One example is the transport of tau protein, which is known for its involvement in Alzheimer’s disease (Scholz and Mandelkow 2014). To be transported from the soma, where it is synthesized, to the axon, tau has to pass the tau diffusion barrier (Li et al. 2011). Tau concentration in the axon is larger than in the soma, thus tau entry into the axon requires transport against the concentration gradient. A model of this process has recently been developed in Kuznetsov and Kuznetsov (2020). Additionally, tau concentration at the axon tip is larger than its concentration in the proximal axon (Black et al. 1996). Thus, in the axon, tau is also transported against its concentration gradient (Kuznetsov and Kuznetsov 2018a, 2018b). Other examples involving anterograde protein transport against the protein’s concentration gradient include dynein, which cannot move in the anterograde direction on its own (Twelvetrees et al. 2016), and synapsin, which is also enriched at synapses (Roy 2020).

## 2. Methods and models

### 2.1. The model of slow (bidirectional) axonal transport of cytosolic proteins

A continuum model of slow axonal transport of neurofilaments (which move in slow component-a (SCa), which is characterized by an average velocity of 0.2-1 mm/day Jensen et al. 1999) was proposed in Jung and Brown (2009). Kuznetsov et al. (2009a, 2009b) extended this model to cytosolic proteins, which are transported in SCb and can diffuse in the cytoplasm. This model extension was done by the addition of diffusion and degradation terms in the free state.

Although our model of slow axonal transport is applicable to any of the proteins transported in SCb, all of which move bidirectionally, here we apply the model to α-syn monomer. α-syn monomer, after being synthesized in the soma, is transported to the presynaptic terminal by slow axonal transport. Slow axonal transport is characterized by short rapid movements, which can be anterograde or retrograde, and are presumably powered by kinesin and dynein motors, respectively. Kinesin and dynein motors run along MTs. Rapid movements are interrupted by prolonged pauses, during which α-syn monomers presumably maintain some association with MTs.

For most of the neuron’s lifetime, the long-range transport processes in the axon operate close to steady state conditions. Therefore, we neglected the time derivatives and simulated axonal transport as quasisteady-state. Since axonal transport is mostly one-dimensional (it occurs along the axon, Fig. 1), we characterized all concentrations in the axon by their linear number densities (number of cargoes in a certain kinetic state per unit length of the axon). The model includes two motor-driven states for α-syn monomers with concentrations 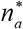 and 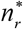, respectively. Conservation of α-syn residing in motor-driven states (Fig. 2a) gives the following equations:

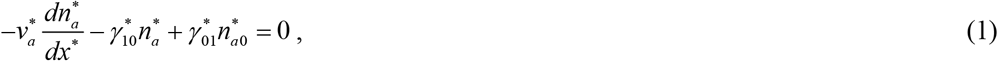

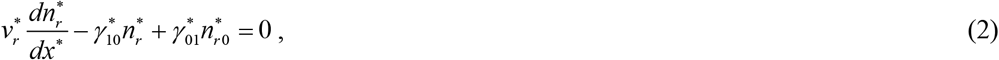

where *x*\* is the Cartesian coordinate along the axon (Fig. 1). Various α-syn concentrations are defined in Table 1 and various model parameters are defined in Tables 2 and 3. In Eqs. (1) and (2) we assumed that the kinetic constants simulating transitions between the anterograde running and pausing states and between retrograde running and pausing states take on the same values. 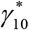 represents the rate of transition to the pausing state and 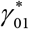 represents the rate of transition to the running state (Fig. 2a).

**Fig. 1.**
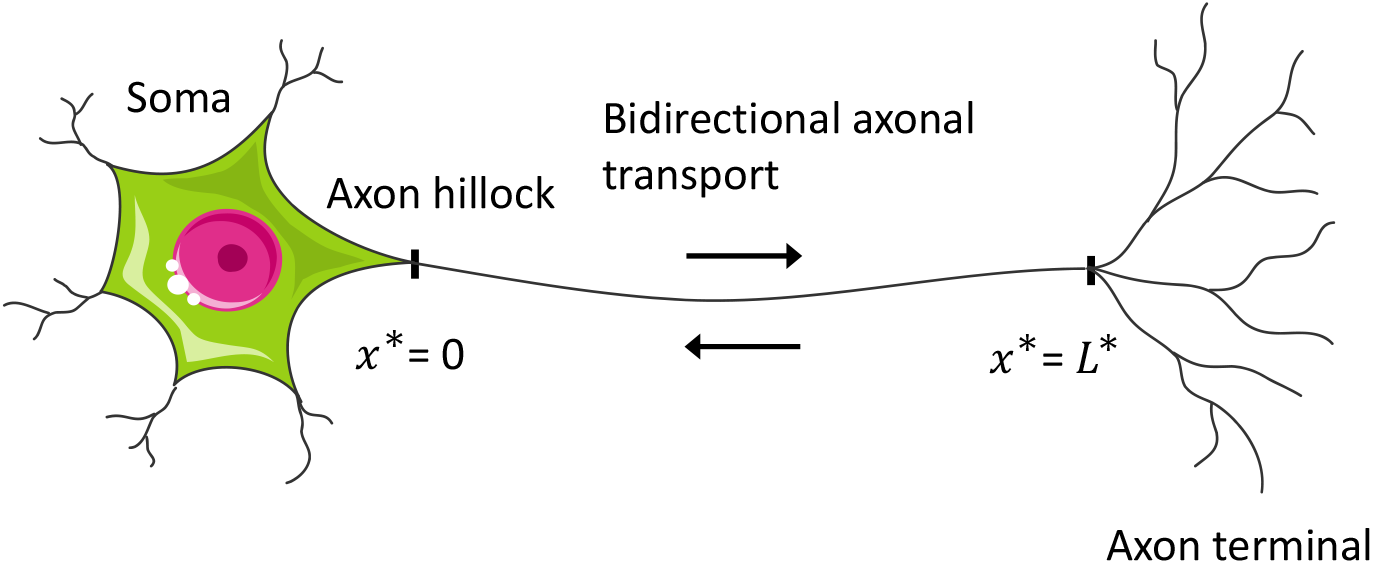
A diagram showing a neuron with an axon and the coordinate system adopted in the model. *Figure generated with the aid of servier medical art, licensed under a creative common attribution 3.0 generic license. http://Smart.servier.com*.

**Table 1.** Dependent variables in the model of α-syn monomers transport from the soma to the presynaptic terminal.

| Symbol | Definition | Units |
| --- | --- | --- |
| $\dot{J}_{tot}^*$ | Total flux of α-syn monomers through a given cross-section in the axon | $s^{-1}$ |
| $n_a^*$ | Concentration of on-track α-syn monomers moving along MTs anterogradely, propelled by molecular motors | $\mu m^{-1}$ |
| $n_r^*$ | Concentration of on-track α-syn monomers moving along MTs retrogradely, propelled by molecular motors | $\mu m^{-1}$ |
| $n_{a0}^*$ | Concentration of pausing on-track α-syn monomers that are still associated with molecular motors and can resume their anterograde motion | $\mu m^{-1}$ |
| $n_{r0}^*$ | Concentration of pausing on-track α-syn monomers that are still associated with molecular motors and can resume their retrograde motion | $\mu m^{-1}$ |
| $n_{free}^*$ | Concentration of free (off-track) α-syn monomers in the cytosol | $\mu m^{-1}$ |

**Table 2.**
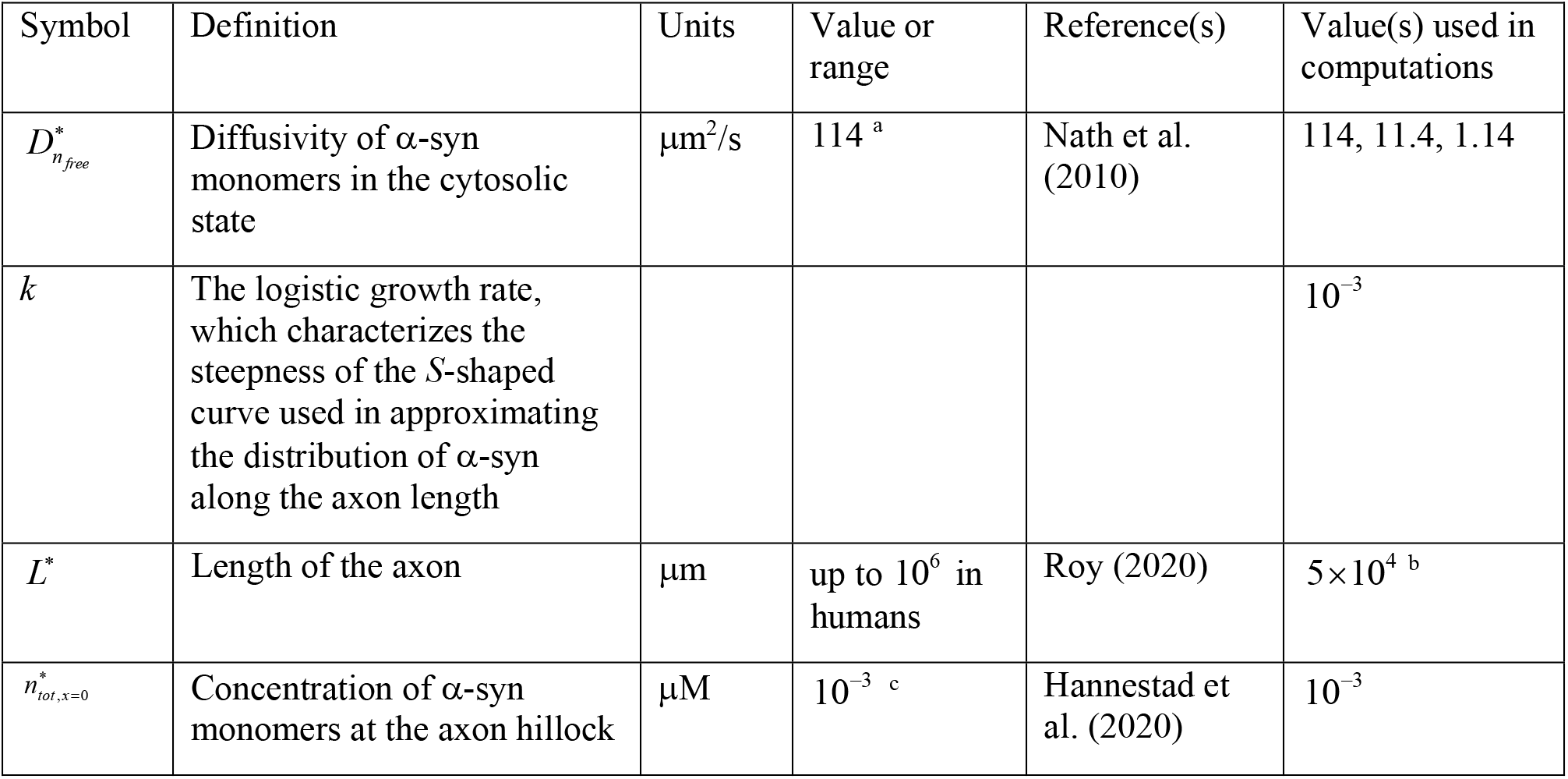
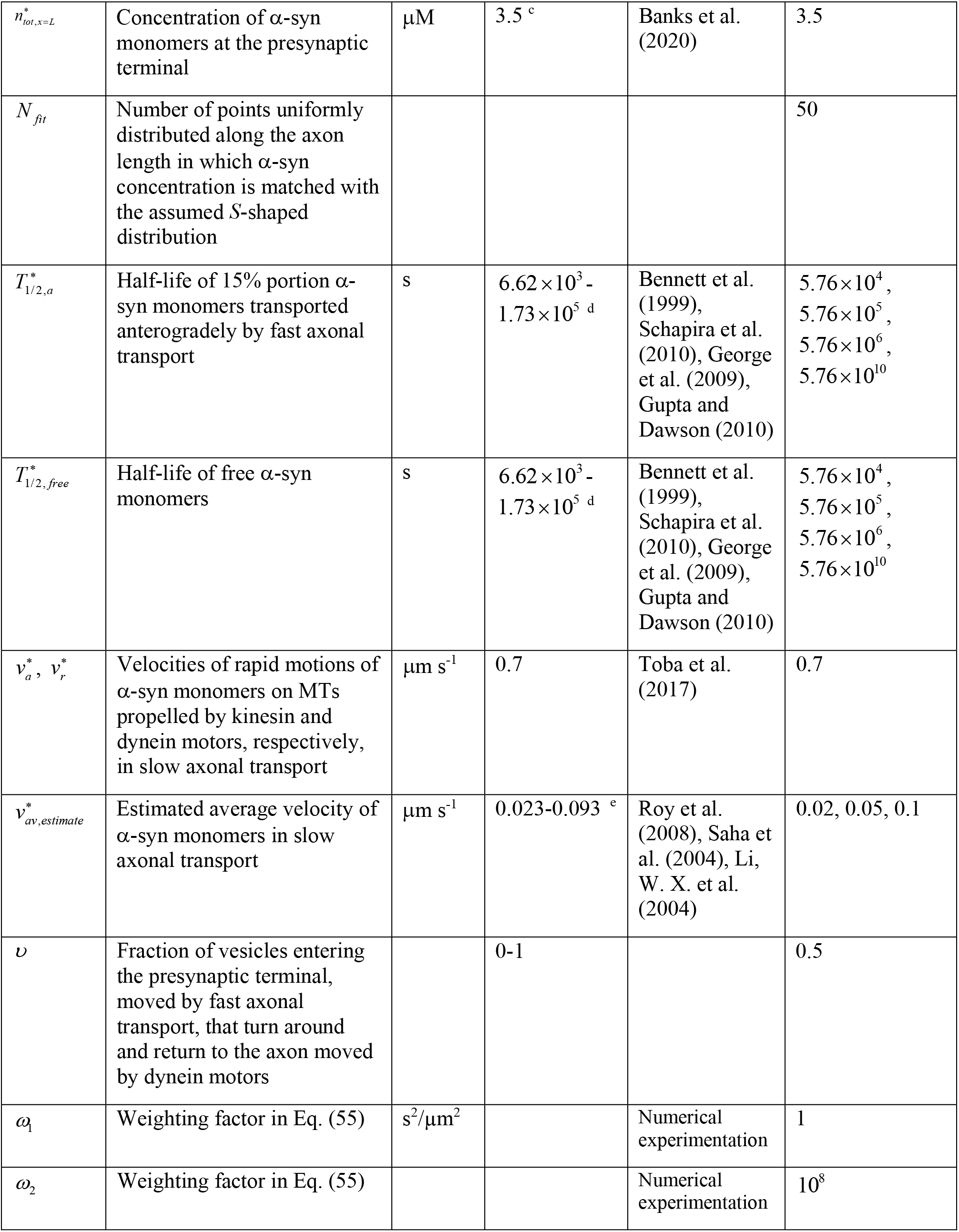
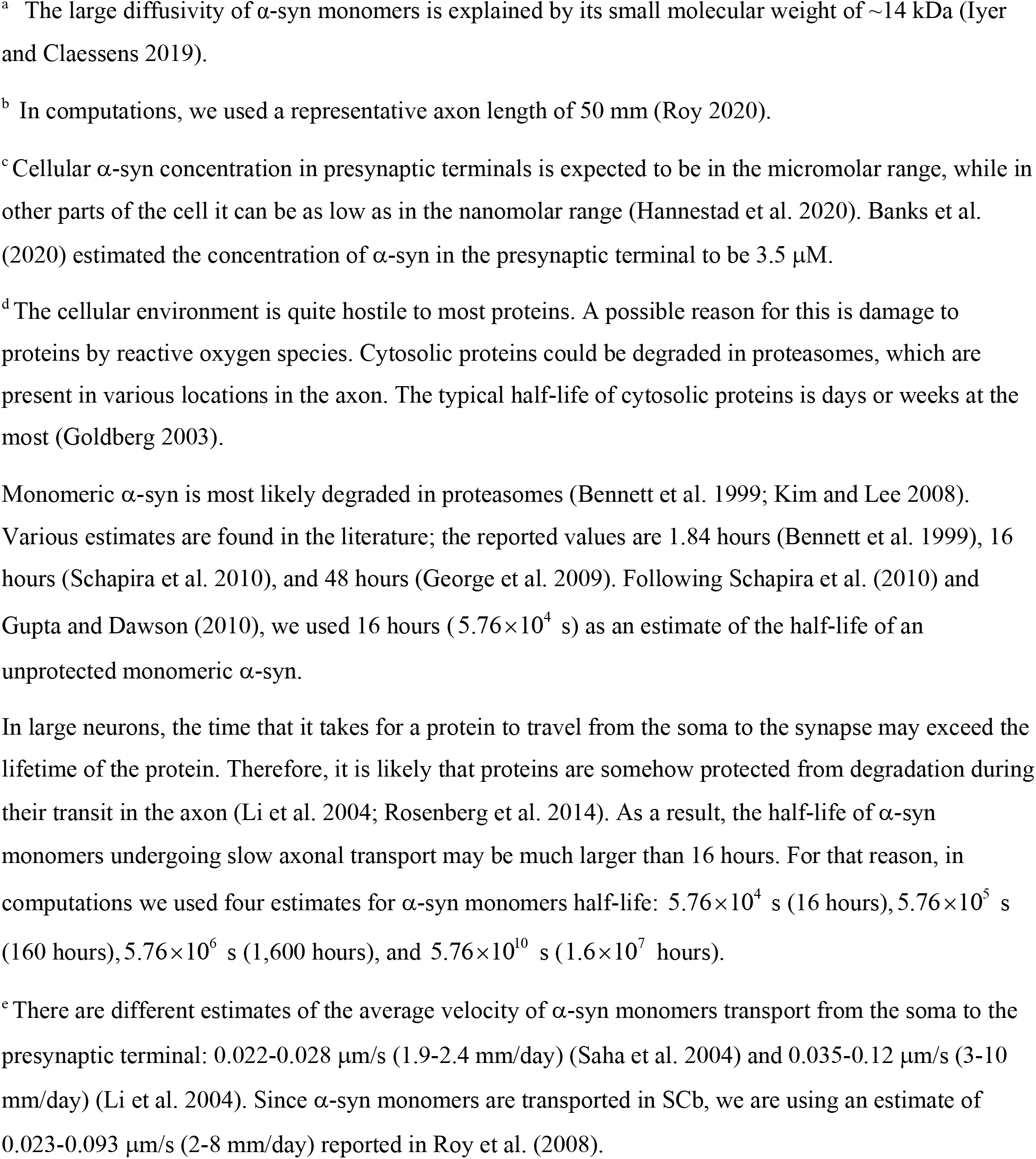
Parameters characterizing transport of α-syn monomers in the axon taken from published data or assumed on physical grounds.

**Table 3.** Kinetic constants characterizing transition of α-syn monomers between different kinetic states in slow axonal transport, which are displayed in Fig. 2a. Values of these kinetic constants were obtained by finding values that give the best fit with respect to different objectives: getting the best fit (i) between the assumed concentration (given by Eq. (56)) and computed α-syn concentration and (ii) between the value of the average α-syn transport velocity reported in the literature (we used three values, 0.02, 0.05, and 0.1 μm/s) and its computed value. These objectives were weighted in defining the objective (penalty) function, see Eq. (55). For optimizing the fit, the Least Square Regression (LSR) was utilized, as described in Kuznetsov and Kuznetsov (2017a, 2017b) and Beck and Arnold (1977).

| Symbol | Definition | Units | Initial guess used in the optimization procedure |
| --- | --- | --- | --- |
| $\gamma_{10}^*$ | Kinetic constant describing the rate of transitions $n_a^* \rightarrow n_{a0}^*$ and $n_r^* \rightarrow n_{r0}^*$ (Fig. 2a) | $\text{s}^{-1}$ | $7.19 \times 10^{-3}$ |
| $\gamma_{01}^*$ | Kinetic constant describing the rate of transitions $n_{a0}^* \rightarrow n_a^*$ and $n_{r0}^* \rightarrow n_r^*$ (Fig. 2a) | $\text{s}^{-1}$ | $4.01 \times 10^{-3}$ |
| $\gamma_{ar}^*$ | Kinetic constant describing the rate of transitions $n_{a0}^* \rightarrow n_{r0}^*$ (Fig. 2a) | $\text{s}^{-1}$ | $4.59 \times 10^{-3}$ |
| $\gamma_{ra}^*$ | Kinetic constant describing the rate of transitions $n_{r0}^* \rightarrow n_{a0}^*$ (Fig. 2a) | $\text{s}^{-1}$ | $6.76 \times 10^{-3}$ |
| $\gamma_{on,a}^*$ | Kinetic constant describing the rate of transitions $n_{free}^* \rightarrow n_{a0}^*$ (Fig. 2a) | $\text{s}^{-1}$ | $9.43 \times 10^{-2}$ |
| $\gamma_{on,r}^*$ | Kinetic constant describing the rate of transitions $n_{free}^* \rightarrow n_{r0}^*$ (Fig. 2a) | $\text{s}^{-1}$ | $5.90 \times 10^{-2}$ |
| $\gamma_{off,a}^*$ | Kinetic constant describing the rate of transitions $n_{a0}^* \rightarrow n_{free}^*$ (Fig. 2a) | $\text{s}^{-1}$ | $5.29 \times 10^{-3}$ |
| $\gamma_{off,r}^*$ | Kinetic constant describing the rate of transitions $n_{r0}^* \rightarrow n_{free}^*$ (Fig. 2a) | $\text{s}^{-1}$ | $5.40 \times 10^{-3}$ |

**Fig. 2.**
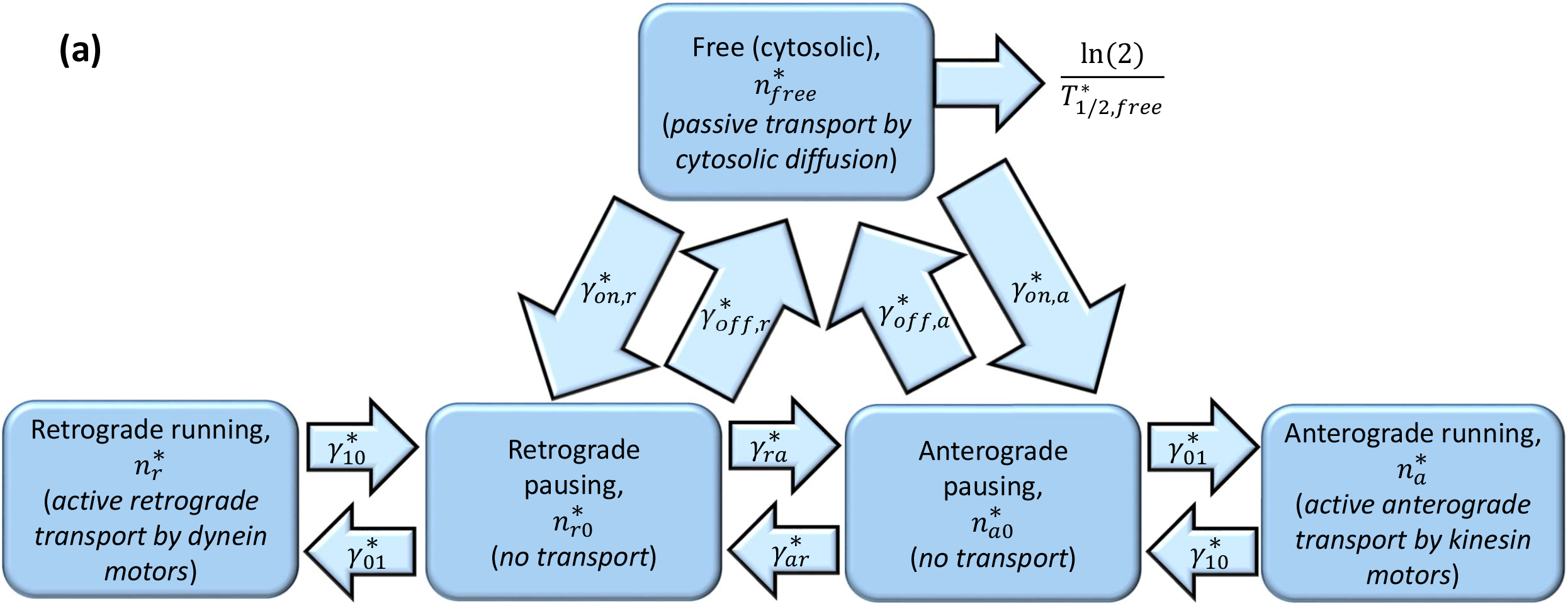
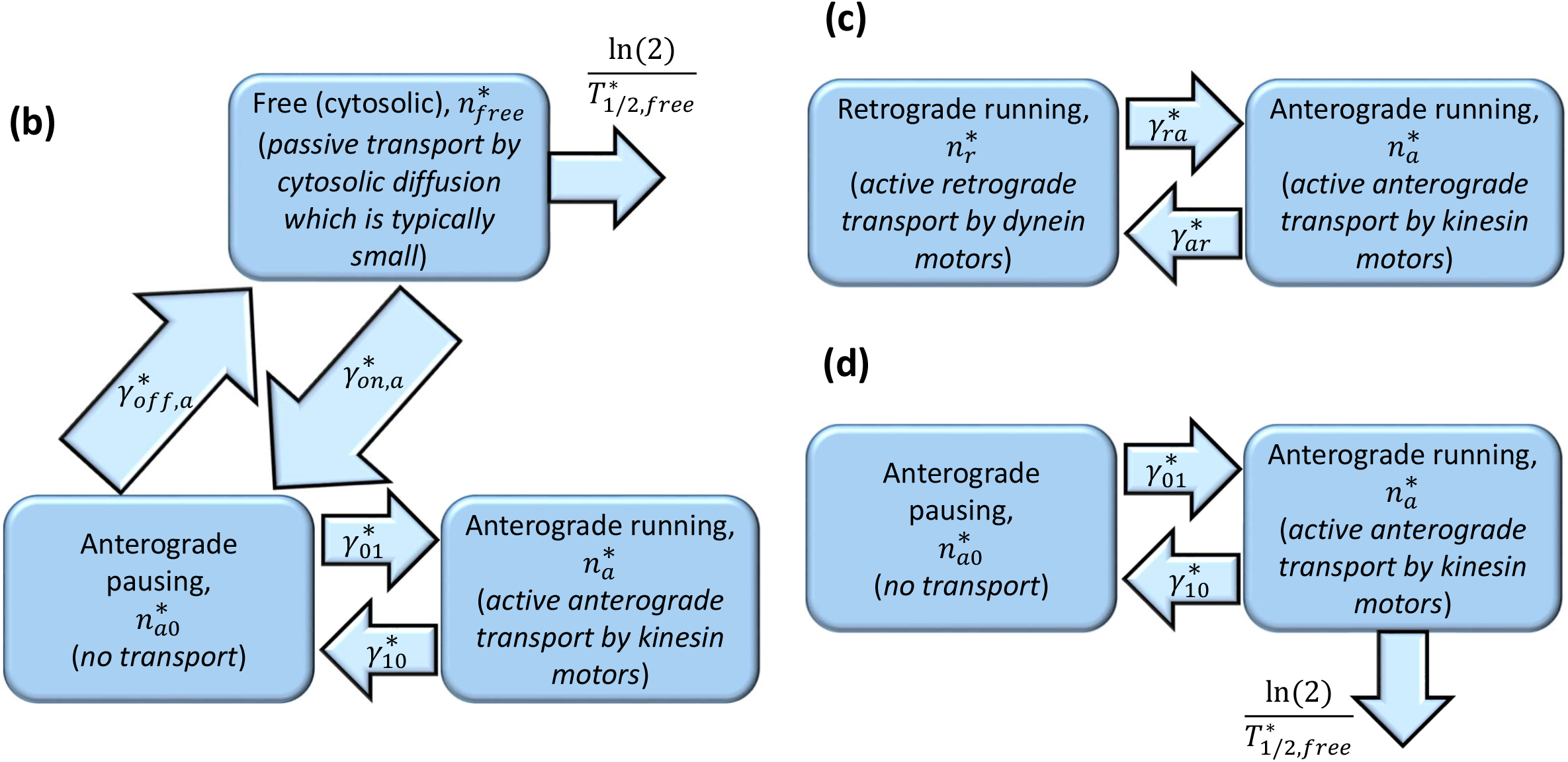
A kinetic diagram showing various kinetic states in our model of α-syn transport in the axon and transitions between these kinetic states. Degradation of free monomeric α-syn due to its destruction in proteasomes is also shown (in the model illustrated in Fig. 2d the degradation process is added in the anterograde state because there is no free state in this model). (a). The diagram is based on the model of SCa transport of neurofilaments developed in Jung and Brown (2009) with modification to this model suggested in Kuznetsov et al. (2009a, 2009b) to extend this model to SCb-transported proteins. Figs. 2b,c,d show various versions of the model with some of the transport modes and kinetic states removed. This is done to investigate what transport modes are minimally required to describe cargo transport against its concentration gradient.

The first terms on the left-hand sides of Eqs. (1) and (2) describe fast motions of α-syn monomers in the anterograde (with a velocity 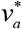) and retrograde (with a velocity 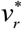) directions, respectively. The other terms describe transitions between motor-driven and pausing kinetic states. Fig. 2a shows various kinetic states and transitions between them described by kinetic rates 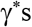.

The model also includes equations for α-syn monomers in two pausing states, with concentrations 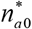 and 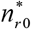, respectively. It is assumed that, despite pausing, α-syn monomers maintain some connection with MTs but are ready to resume anterograde or retrograde motion. Equations expressing conservation of α-syn monomers in the pausing states involve only terms that describe transitions to/from other kinetic states, since no motor-driven transport or diffusion occurs in the pausing states (Fig. 2a):

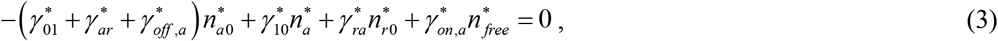

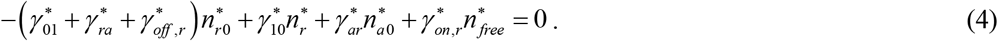

SCb-transported proteins have diffusible fractions (Roy 2014). Free (cytosolic) α-syn monomers can thus be transported by diffusion. They also can be destroyed by degradation in proteasomes (Raichur et al. 2006), which are present in various locations in the axon. Free α-syn can also transition to anterograde or retrograde biased pausing states (Fig. 2a). Stating the conservation of free monomeric α-syn gives the following equation:

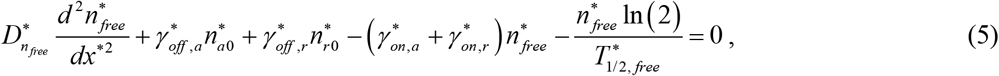

where 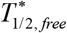 is the half-life of free α-syn monomers. The last term on the left-hand side of Eq. (5) accounts for the destruction of free α-syn monomers in proteasomes. In this term, the rate of decay of free α-syn is assumed to be directly proportional to the concentration of the free α-syn.

Eqs. (1)–(5) must be solved subject to the following boundary conditions:

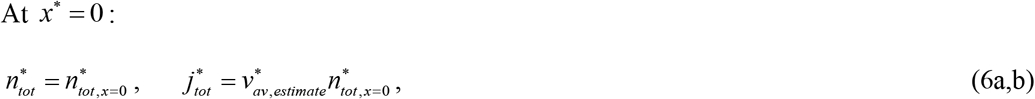

where 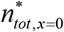 is the total concentration of α-syn monomers at the axon hillock (at *x*=0), which is assumed to be constant. We estimated its value in Table 2.

In our model, all *γ*\* s in Eqs. (1)–(5) are constants that are independent of the cargo concentration along the axon.

Eq. (6a) postulates the total concentration of α-syn monomers at the axon hillock and Eq. (6b) postulates the total flux of newly synthesized α-syn monomers entering the axon. In Eq. (6b)

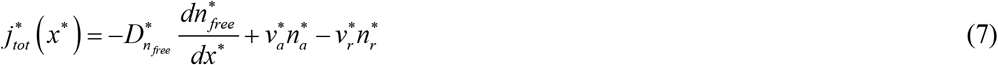

is the total flux of α-syn monomers due to diffusion in the cytosol (the first term of the right-hand side of Eq. (7)) and motor-driven anterograde (the second term) and retrograde (the third term) transport.

The total concentration of α-syn monomers (in all five kinetic states displayed in Fig. 2a) is found as:

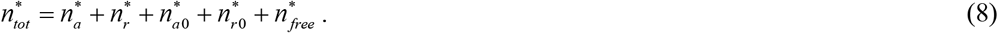

As we suggested in Kuznetsov and Kuznetsov (2015), the average velocity of a protein in slow axonal transport is defined as the ratio of the total protein flux to the total protein concentration:

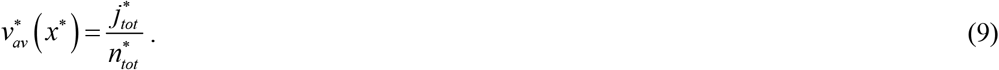

At the axon presynaptic terminal, we imposed the following boundary conditions:

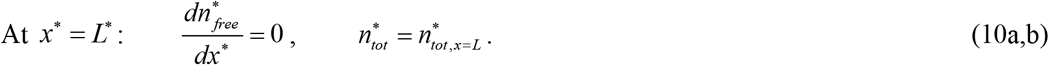

We defined the dimensionless concentration of α-syn as:

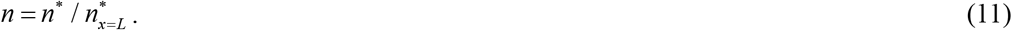

This equation applies to the total α-syn concentration and all α-syn concentration components (*n_a_*, *n_r_*, etc.). Eq. (10b) can then be recast as:

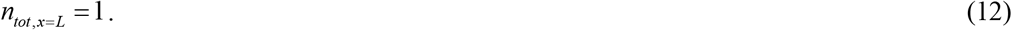

The dimensionless total flux of α-syn is defined as:

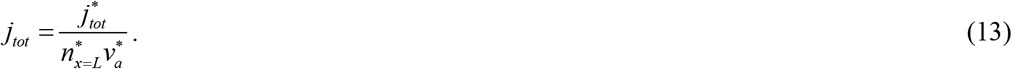

### 2.2. The model of fast anterograde (unidirectional) axonal transport of various organelles

Fast anterograde axonal transport of various membranous organelles is modeled by the following equation:

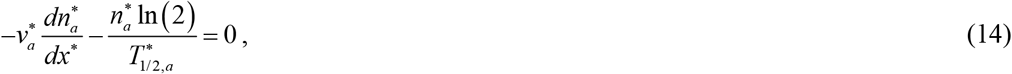

where 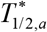 is the half-life of cargos transported by fast axonal transport. In Eq. (14) we neglected the vesicles’ diffusivity due to their large size. This equation must be solved subject to the following boundary condition:

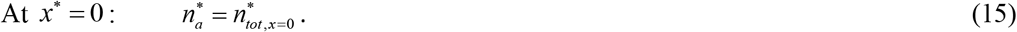

To enable a comparison between slow and fast axonal transport simulations, we assumed the same protein concentration at *x*\* = 0 for fast axonal transport as for slow axonal transport (compare with Eq. (6a)).

The analytical solution of Eq. (14) with boundary condition (15) is

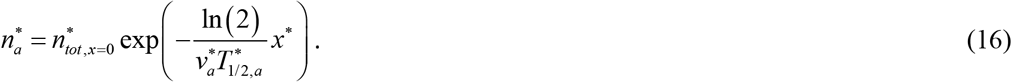

Eq. (16) applies to the whole length of the axon, 0 ≤ *x*\* ≤ *L*\*.

Wong et al. (2012) has shown that neuropeptide delivery to en passant boutons in type Ib terminals of Drosophila motoneurons involves dense core vesicle (DCV) circulation, in which DCVs that reached the most distal bouton in the terminal turn around and move back by retrograde transport. The concentration of retrogradely transported cargos can be modeled by the following equation:

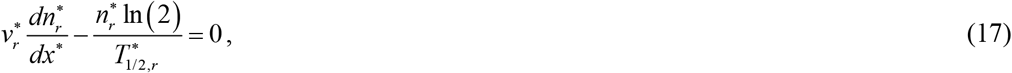

where 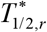 is the half-life of cargos transported by retrograde transport.

This equation must be solved subject to the following boundary condition at the axon tip:

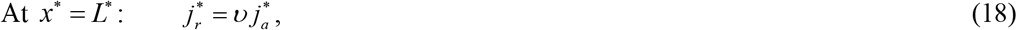

where 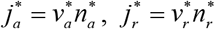, and *υ* is the fraction of vesicles entering the presynaptic terminal that turn around and return to the axon, moved by dynein motors. Under steady state conditions, the difference, 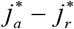, must be destroyed in the presynaptic terminal (or released from it). Eq. (18) can be recast as:

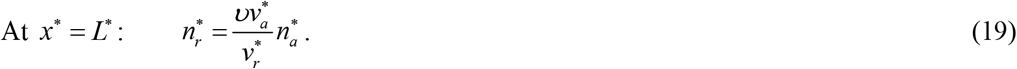

If 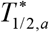 is very large, then, from Eq. (16), 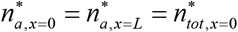, and the solution of Eq. (17) with boundary condition (19) is given by the following equation:

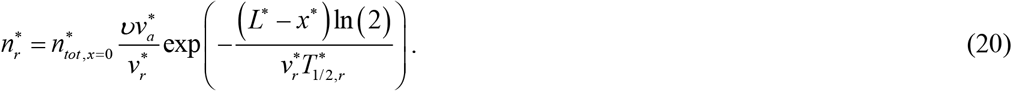

Similar to Eq. (16), Eq. (20) applies for 0 ≤ *x*\* ≤ *L*\*.

### 2.3. The model of slow axonal transport of cytosolic proteins with 15% fast component

According to Jensen et al. (1999), Utton et al. (2005), Tang et al. (2012), and Roy (2014), 10-15% of proteins transported in SCb are moved by fast axonal transport. In order to describe this situation, we utilized the above models of fast and slow axonal transport, in which Eq. (6a,b) was modified as:

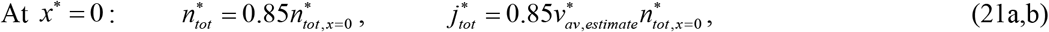

and Eq. (15) is replaced with the following equation:

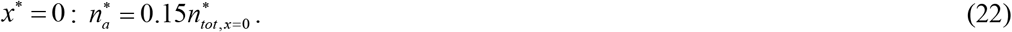

### 2.4. Axonal transport model that includes anterograde motor-driven transport with pausing and diffusion only

#### 2.4.1. General formulation of the anterograde-diffusion cargo transport model

In this section, we investigate the question of whether an anterograde transport model with pausing and diffusion in the free state can describe transport against the concentration gradient (Fig. 2b). Consider cargo assembled in the cell body. It binds to kinesin which transports it along the MTs in the axon in the anterograde direction towards the terminal. The cargo may detach during its journey and diffuse along the axon before it binds again to kinesin to continue its journey to the axon terminal. After arriving at the nerve terminal, the cargo may detach from its kinesin motor and accumulate and degrade, building up a gradient similar to that shown in Fig. 4a. In this case, the mathematical model given by Eqs. (1)–(5) reduces to the following equations:

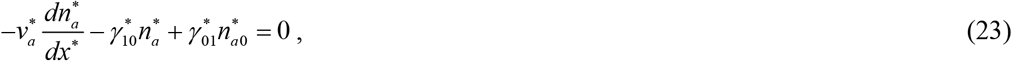

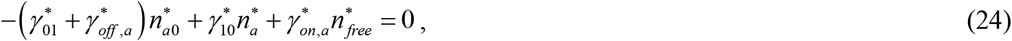

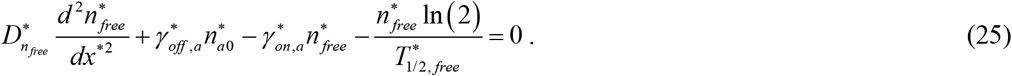

Eqs. (23)–(25) must be solved subject to the following boundary conditions:

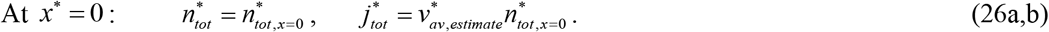

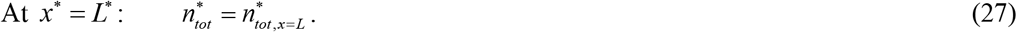

#### 2.4.2. A perturbation solution of the anterograde-diffusion cargo transport model for the case of small cargo diffusivity

Cargos transported by slow axonal transport are usually assembled in some kinds of complexes. Neurofilaments, which move in SCa, are assembled as polymers (Yan and Brown 2005; Maday et al. 2014). Cytosolic proteins, such as synapsin, assemble in multiprotein complexes before being moved in SCb (Scott et al. 2011; Maday et al. 2014). Studies of radiolabeled wave-profiles of various SCb proteins suggest that they organize themselves in multicargo complexes, and different SCb proteins are transported together (Roy et al. 2007; Roy et al. 2008; Roy 2014). The diffusivity of such complexes is expected to be extremely small.

Motivated by this, we used the perturbation technique to investigate the limit of the anterograde-diffusion transport model given by Eqs (23)–(25) for the case when cargo diffusivity approaches zero. The question was whether in this model a cargo can be transported against its concentration gradient.

Using Eq. (24), the following is obtained:

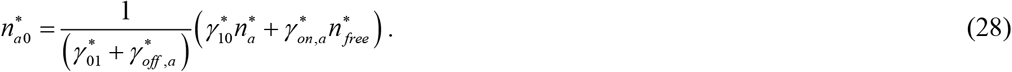

Eq. (28) is used to eliminate 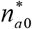 from Eq. (23), which becomes:

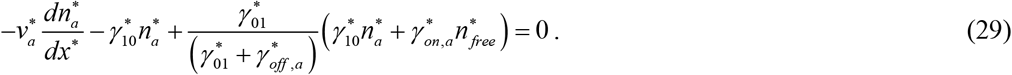

Eliminating 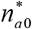 from Eq. (25), this equation becomes:

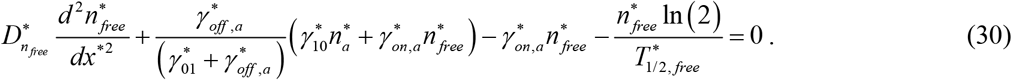

We assumed that 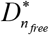 is a small parameter. The following perturbation expansions are utilized:

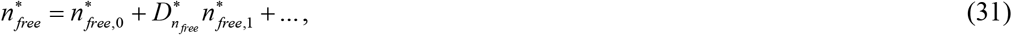

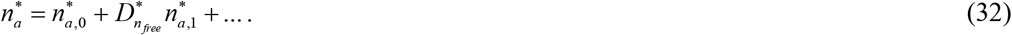

By substituting Eq. (32) into Eq. (29), separating the terms that do and do not contain the small parameter 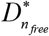, and equating the terms that do not contain 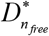 to zero, the following is obtained:

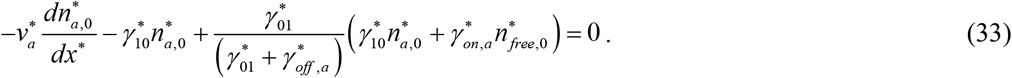

By substituting Eqs. (31) and (32) into Eq. (30), separating the terms that do and do not contain the small parameter 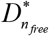, and equating the terms that do not contain 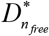 to zero, the following is obtained:

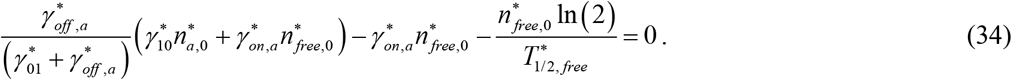

Solving Eq. (34) for 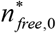, the following is obtained:

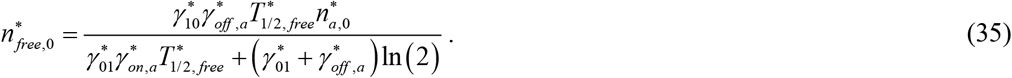

Substituting Eq. (35) into Eq. (33) to eliminate 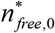, the following is obtained:

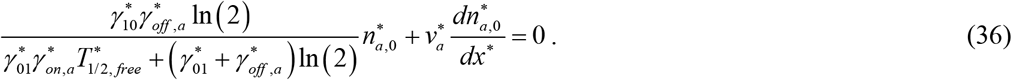

For the case when 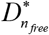 is small and there is no retrograde motor-driven transport, the flux of cargos is given by

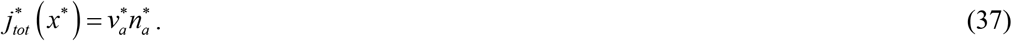

Eq. (36) is solved subject to the following boundary condition:

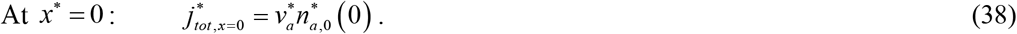

The solution for 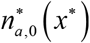 is

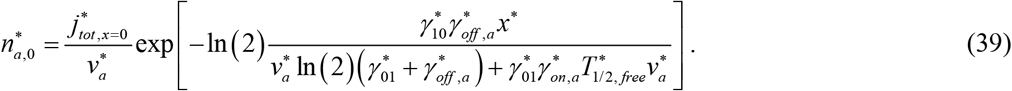

The total cargo concentration can then be obtained using Eqs. (28), (35), and (39) as:

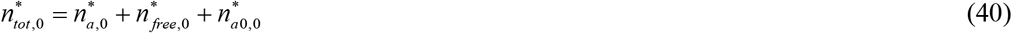

### 2.5. Axonal transport model that includes anterograde and retrograde motor-driven transport without diffusion

In this section we investigate the situation when diffusion and pausing states are dropped from the model described in section 2.1, and only motor-driven states are retained (Fig. 2c). Eqs. (1)–(5) in this case reduce to:

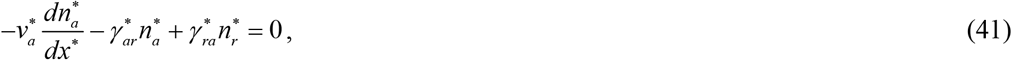

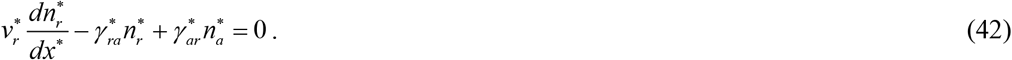

Eqs. (41) and (42) must be solved subject to the following boundary conditions:

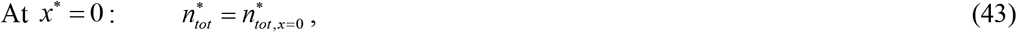

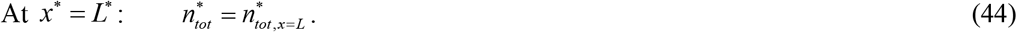

The total flux of cargo in this case is calculated as:

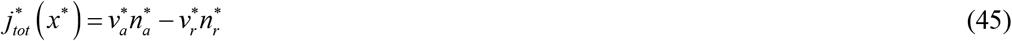

and the total cargo concentration is found as:

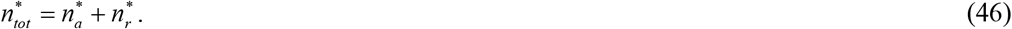

### 2.6. Axonal transport model that includes anterograde transport with pausing only

In this section we investigate what happens if diffusion is dropped out of the model presented in section 2.4, and whether the model, in this situation, can still correctly predict transport against the concentration gradient (Fig. 2d). Eqs. (23)–(25) in this case reduce to:

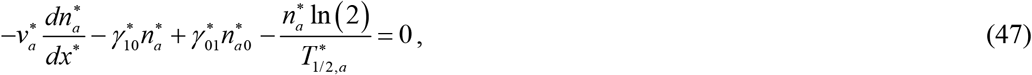

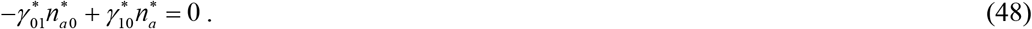

Similar to Eq. (14), we added a term in Eq. (47), which describes a possible cargo decay in the motor-driven state. Without this term, the equations exhibit a trivial (constant concentration) solution.

Eqs. (47) and (48) must be solved subject to the following boundary condition:

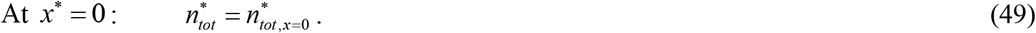

By adding Eqs (47) and (48), Eq. (14) is obtained. The solution of Eq. (14) in this case is

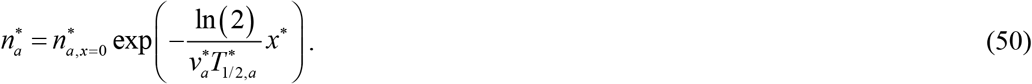

Using Eq. (48) and 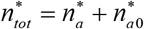 results in

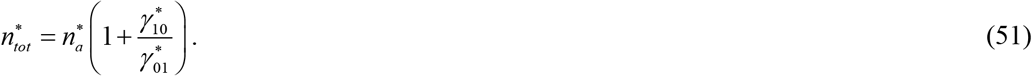

Utilizing Eq. (50), Eq. (51) is recast as:

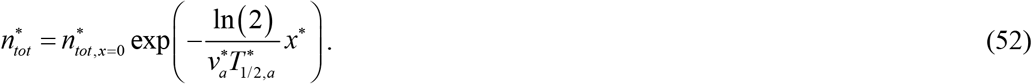

Eq. (52) means that anterograde transport with pausing cannot capture transport against the concentration gradient correctly. Note that modulating values of kinetic constants 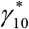 and 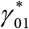 (allowing them to depend on *x*\*) cannot fix this problem. The issue with this model is that it describes cargo transport in one (anterograde) direction only, and thus it cannot account for cargo transport in the retrograde direction. In order to correctly describe the physical situation (a given high cargo concentration at the axon tip), the model must allow for the imposition of the second boundary condition at the tip, which is not possible because the highest order derivative in the governing equations is of the first order (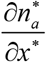, Eq. (47)).

The analysis above shows that the case with cargos which are anterogradely transported by motors and can also pause is reduced to the case of anterogradely motor-driven cargos. The reason why the unidirectional anterograde motor-driven transport model cannot describe cargo transport against the concentration gradient is further explained in Fig. 3. Under steady state-conditions (due to a long duration of human life, most of the time axonal transport can be assumed to operate at steady state), the balance of cargos entering and leaving a control volume (CV) can be stated as follows:

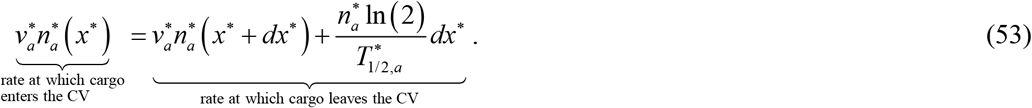

**Fig. 3.**
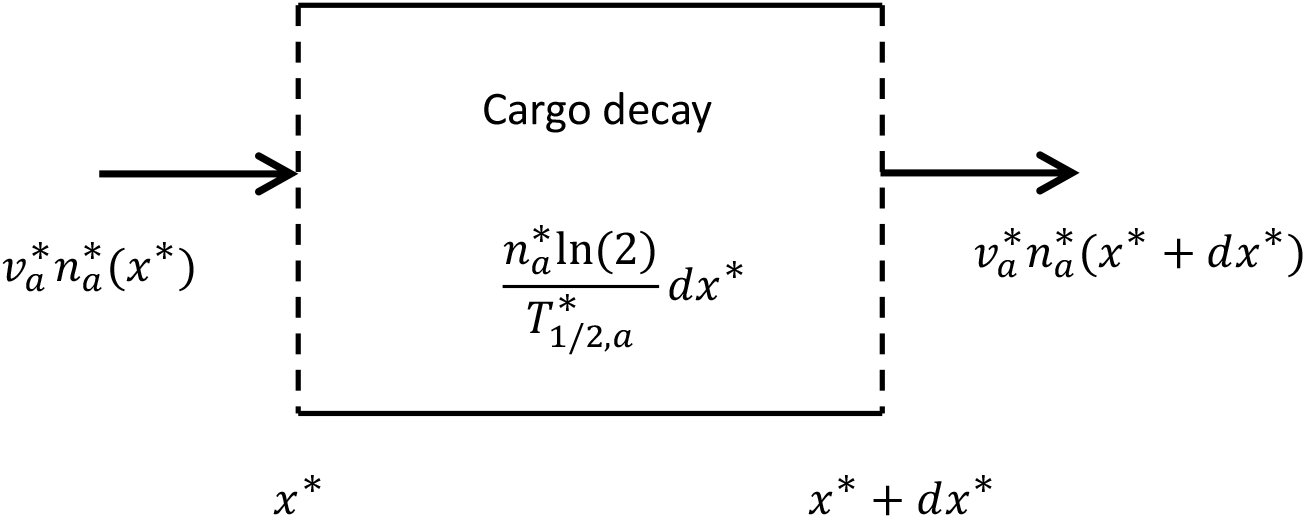
A schematic diagram showing a control volume (CV) in the axon. Anterogradely transported cargo can enter and leave the CV; the cargo can also decay in the CV.

From Eq. (53) it follows that

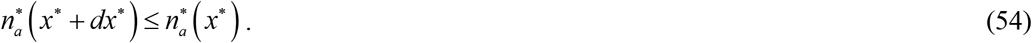

Equality is for the case of 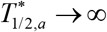.

### 2.7. Finding values of kinetic constants by least square regression

In order to solve the problem numerically, values of eight kinetic constants given in Table 3 are needed. These values can be found by using multi-objective optimization (see, for example, Kool et al. 1987; Zadeh 2008; Zadeh and Shah 2010; Zadeh 2011; Zadeh and Montas 2014). The following objective (penalty) function was minimized:

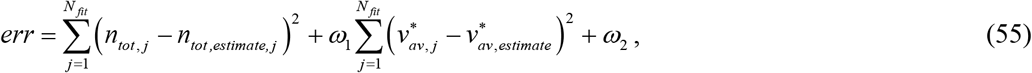

where the axon was discretized into *N_fit_* uniformly spaced points in which cargo concentration and velocity were matched with the synthetic data.

For the most general slow axonal transport model formulated in section 2.1, the objective function defined by Eq. (55) depends on eight parameters: 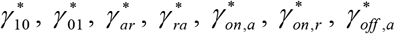, and 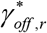. The best-fit values of these parameters were determined by finding a set that gives the minimum value of the objective function.

The objective function defined by Eq. (55) combines three different effects. The first term on the right-hand side of Eq. (55) estimates how well the α-syn concentration predicted by the model, *n_tot, j_*, fits the estimated concentration of α-syn given by the following sigmoid curve, which we obtained by modifying a logistic function:

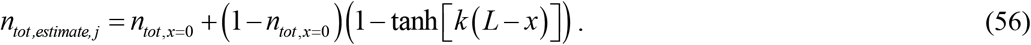

Although synthetic data given by Eq. (56) represent an α-syn monomer distribution that would require α-syn transport against the concentration gradient, this gradient is not uniform along the axon length; most of the increase is assumed to occur near the presynaptic terminal. This is done to model the presynaptic localization of α-syn monomers.

The second term on the right-hand side of Eq. (55) estimates how well the average velocity of α-syn predicted by the model, 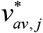, fits the estimated average α-syn velocity, 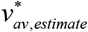. We used three values for 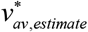, which are in the range of average velocities of SCb (2-8 mm/day, Roy et al. 2008): 0.02, 0.05, and 0.1 μm/s. We also used *ω*_1_=1 s^2^/μm^2^. This value was selected by numerical experimentation to avoid overfit in terms of either α-syn concentration or average velocity, as described in Kuznetsov and Kuznetsov (2017b).

Physically, α-syn concentrations in various kinetic states must be non-negative. Therefore, we set *ω*_2_ = 10^8^ if any of *n_a,j_*, *n_r, j_*, *n*_*a*0,*j*_, *n*_*r*0,*j*_, and *n_free, j_* (*j*=1, …, *N_fit_*) are negative. If the half-life of α-syn monomers is infinitely large, its flux under steady state conditions must stay constant. If the half-life of α-syn is finite, its flux decreases as *x\** increases. However, since the presynaptic terminal must be supplied with α-syn monomers, physically the flux must remain positive. Therefore, we also set *ω*_2_ = 10^8^ if *j_tot_* (*j*=1, …, *N_fit_*) becomes negative.

Examples of the best fit-values of model parameters for models presented in sections 2.1, 2.4, and 2.5 are given in Tables S1, S2, and S3, respectively.

### 2.7. Numerical solution

We eliminated 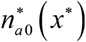 and 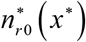 from Eqs. (1)–(5) using Eqs. (3) and (4). This led to a system of ordinary differential equations of the fourth order, which was solved using MATLAB’s BVP5C solver (MATLAB R2019a, MathWorks, Natick, MA, USA). The inverse problem, which involves a determination of values of kinetic constants (Table 3), was solved using MultiStart with the local solver fmincon; these routines are included in MATLAB’s Optimization Toolbox. 1000 random points plus the initial point given in Table 3 were utilized as starting points in searching for the global minimum. The numerical procedure is described in detail in Kuznetsov and Kuznetsov (2018b).

## 3. Results

### 3.1. Bidirectional model can describe transport against the concentration gradient for different average velocities of α-syn monomers

A bidirectional transport model (in particular, a slow axonal transport model) can fit a protein distribution characterized by a uniform low protein concentration in the axon and a sharp concentration increase at the presynaptic terminal. For the simulated case of slow axonal transport of α-syn, the fitting is done by adjusting the values of the kinetic constants characterizing protein transition rates between different kinetic states. The best-fit values of the rate constants for 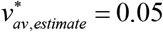 μm s^−1^ are given in Table S1. The model can fit a distribution for any average velocity of α-syn transport within the SCb range (the total concentration is shown in Fig. 4a, the components of the total concentration are shown in Figs. S1–S3). This is because the bidirectional axonal transport model is described by a higher order system of differential equations (second when diffusion of free proteins in the cytosol is neglected and fourth when it is accounted for), and thus allows for the imposition of boundary conditions at the axon hillock and at the presynaptic terminal. Because the model includes both anterogradely and retrogradely transported protein populations, cargo in this model can be transported in both anterograde and retrograde directions, which allows a bidirectional axonal transport model to account for the imposed protein concentration at the presynaptic terminal. This enables bidirectional axonal transport to move proteins against their concentration gradient, unlike pure diffusion (without any other kinetic states but the free state) or unidirectional fast axonal transport.

**Fig. 4.**
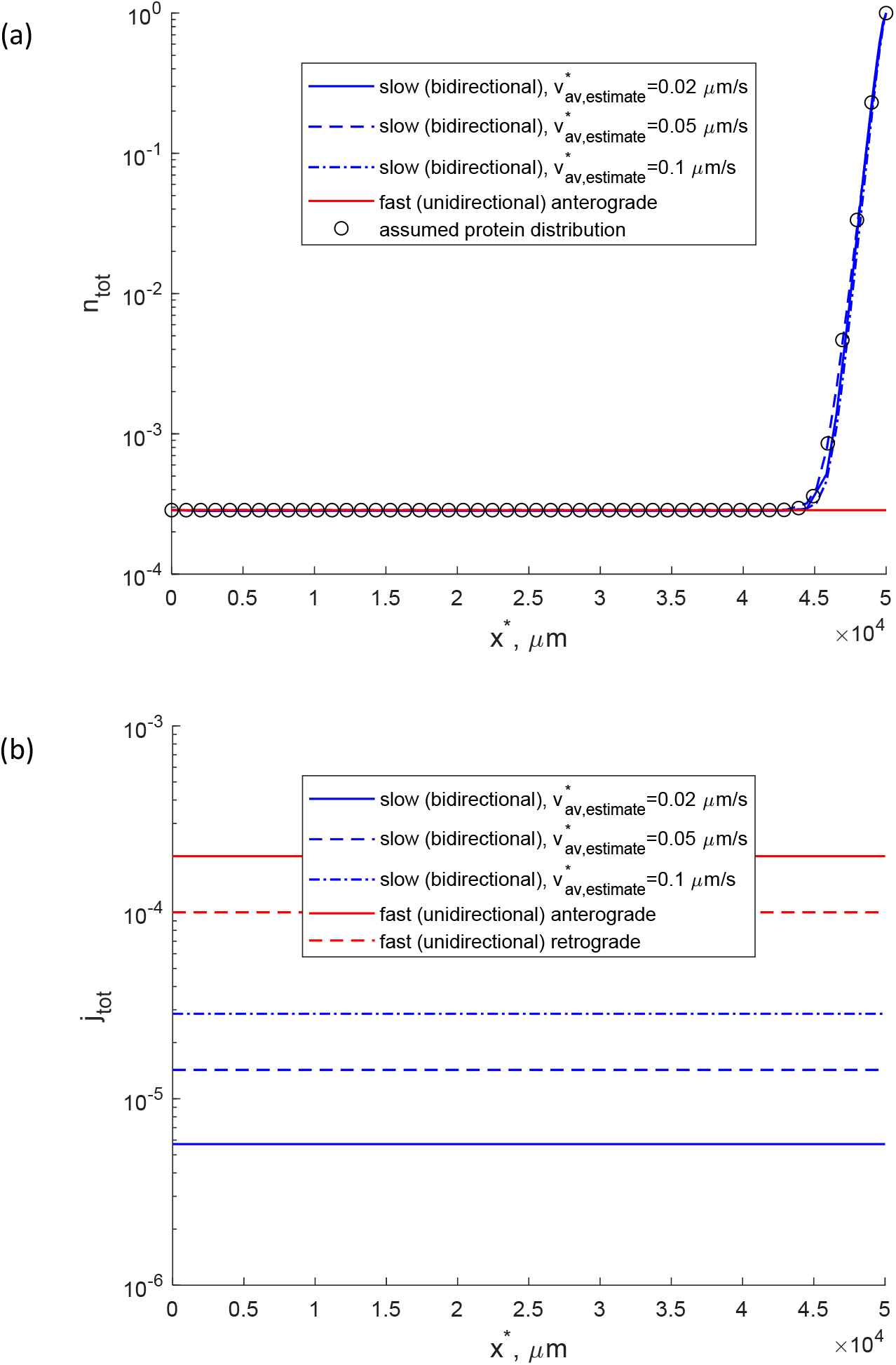
Slow (bidirectional) and fast (unidirectional) axonal transport. 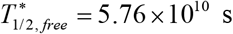, the fraction of α-syn monomers conveyed in the fast component of axonal transport is assumed to be zero. (a) Total concentration of α-syn monomers (the sum of α-syn concentrations in motor-driven, pausing, and diffusing states). An exponential increase in the concentration of α-syn monomers near the tip signifies that α-syn monomers are predominantly localized in the presynaptic terminal. (b) Total flux of α-syn monomers due to the action of molecular motors and diffusion in the cytosol.

Unlike the bidirectional axonal transport model, the unidirectional axonal transport model cannot describe the increase of protein concentration near the presynaptic terminal. If the protein half-life is infinitely large, the protein concentration is uniform and constant (Fig. 4a). This is explained by the fact that the system of governing equations modeling unidirectional (in this case, anterograde) axonal transport is of the first order, and thus can only include one boundary condition, which in the case of anterograde transport has to be imposed at the axon hillock. This is because the model of unidirectional fast anterograde axonal transport includes only transport by anterograde motors, and thus cargo can only be transported in the anterograde direction. Diffusion is neglected in the model of unidirectional transport because this mode of transport usually applies to fast axonal transport, in which case cargos are moved inside vesicles, which are usually too large to exhibit a significant diffusivity.

The slow axonal transport model can describe various cargo distributions. Possible distributions are not limited by a distribution with a constant concentration of cargo along most of the axon, which is followed by a gradient at the terminal (as in the curves shown in Fig. 4a). The distribution in Fig. 4a is a consequence of using a modified logistic function, given by Eq. (56), to represent synthetic data to fit our model. For example, Fig. 3(b) in Kuznetsov and Kuznetsov (2018b) and Fig. 3(a) in Kuznetsov and Kuznetsov (2020) show that the slow axonal transport model can fit a cargo distribution that changes along the whole length of the axon, not only near the tip.

In some situations, cargos participating in fast axonal transport can turn around at the end of the axon and move backward by retrograde transport, propelled by cytoplasmic dynein. This situation may lead to cargo circulation (Wong et al. 2012). However, without cargo transitions between the anterograde and retrograde states, this situation does not allow for the imposition of a higher cargo concentration at the end of the axon. This is because only a fraction of anterogradely transported cargos can turn around, and due to the conservation of cargos this fraction cannot exceed unity. The red dashed curve in Fig. 4b shows the absolute value of the retrograde flux for the situation when half of the anterogradely transported cargos turn around at the end of the axon. The velocities of fast anterograde and retrograde axonal transport components are the same (Fig. 5a).

**Fig. 5.**
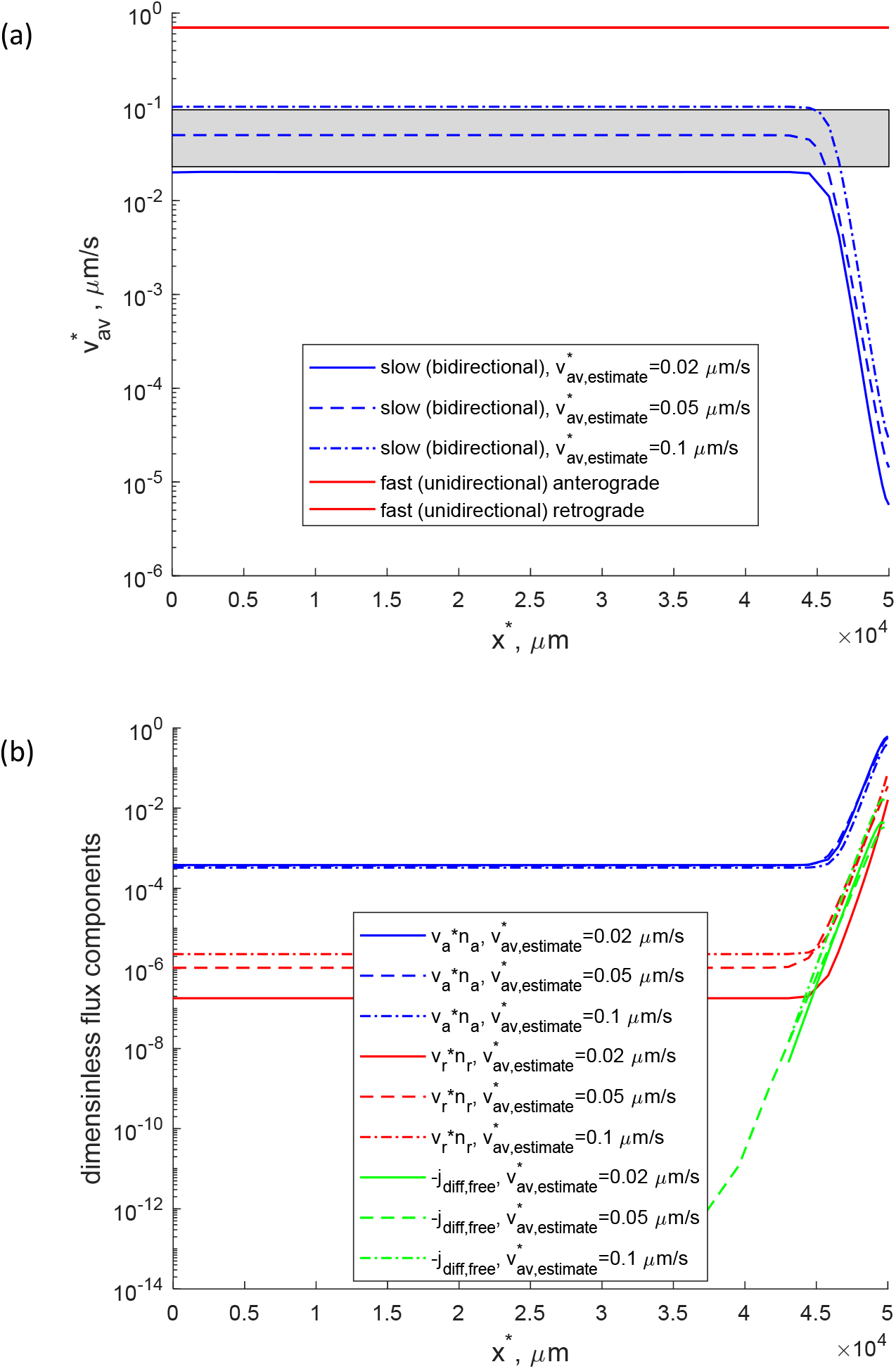
Slow (bidirectional) and fast (unidirectional) axonal transport. 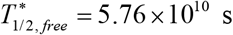, the fraction of α-syn monomers conveyed in the fast component of axonal transport is assumed to be zero. (a) Average velocity of α-syn monomers in slow axonal transport along the axon. (b) Anterograde motor-driven, retrograde motor-driven, and diffusion-driven components of the total α-syn flux. Note that because of the log scale used, Fig. 5b shows the absolute values of the flux components. Since retrograde and diffusion-driven flux components are negative, Fig. 5b shows the negative of these components.

Our model shows that α-syn flux along the axon is constant and uniform if α-syn destruction during its axonal transport is neglected. Since α-syn concentration at the entrance to the axon is assumed to be constant, the constant value of α-syn flux increases with an increase of the average α-syn velocity. For the same α-syn concentration at the axon entrance, fast axonal transport can deliver a much larger α-syn flux due to larger average velocity (Fig. 4b).

The velocity along most of the axon length remains constant, but it decreases close to the presynaptic terminal (Fig. 5a) in the region where the average α-syn concentration increases (Fig. 4a). The increase of α-syn concentration near the axon tip is explained by the increase of both anterograde and retrograde components of axonal transport; the diffusion component remains relatively small (Fig. 5b).

In Supplementary Materials we analyzed the situation with various half-lives of α-syn monomers (section S2.2) and the situation when 15% fraction of α-syn monomers is moved by fast axonal transport (section S2.3). The results show that these situations can also be successfully described by the bidirectional transport model.

### 3.2. The model without retrograde motor-driven transport fails to describe transport against cargo concentration gradient if cargo diffusivity is small

Since proteins are usually transported as multi-protein complexes in slow axonal transport (Puthanveettil et al. 2008; Roy et al. 2008), the investigation of the effect of cargo diffusivity on slow axonal transport is pertinent. To show the effect of cargo diffusivity on the full slow axonal transport model, given by Eqs. (1)–(5), we utilized best-fit values of kinetic constants given in Table S1 in Supplemental Materials. These values minimize the objective function given by Eq. (55) for the case of 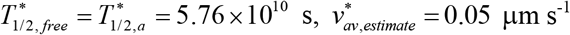, and 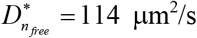. When diffusivity was changed (for example in Fig. 6), the kinetic constants were not changed. This was done to show the effect of cargo diffusivity (otherwise, by modifying values of kinetic constants, a perfect fit with synthetic data shown by hollow circles in Fig. 6 could be obtained). For the case of the full slow axonal transport model, the total concentration of α-syn monomers is only weakly dependent on the diffusivity of α-syn monomers (Fig. 6). This is because the main contributors to the total flux of α-syn monomers are the motor-driven fluxes (anterograde, driven by kinesin, and retrograde, driven by dynein motors) rather than the diffusion-driven flux of α-syn monomers.

**Fig. 6.**
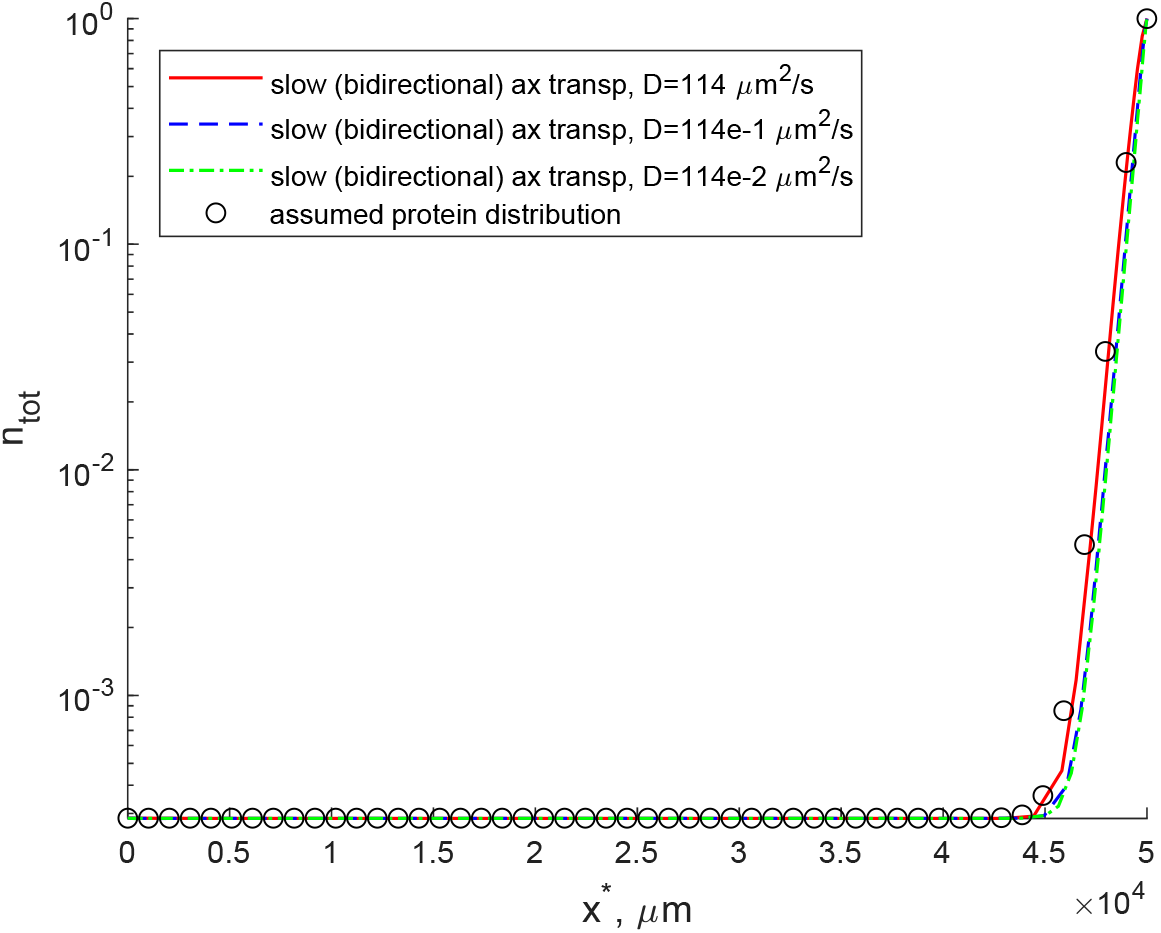
The effect of cargo diffusivity on the total concentration of α-syn monomers computed utilizing the full slow axonal transport model, which includes anterograde, retrograde, and diffusion transport of α-syn monomers. The MATLAB notation is utilized, i.e., 114e-2 means 114×10^−2^. 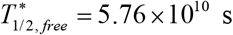 and 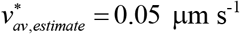.

Next, we investigated the effect of cargo diffusivity on a model that accounts only for anterograde and diffusion-driven cargo transport (with pausing states), which is given by Eqs. (23)–(25). In this case values of kinetic constants are given in Table S2. These were again determined for 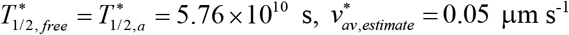, and 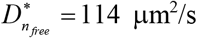, but the number of kinetic constants in this model is smaller than the number of kinetic constants in the full slow axonal transport model (Table S1).

Anterograde (*n_a_*), free (*n_free_*), and pausing (*n*_*a*0_) components of the total α-syn monomer concentration (*n_tot_*) obtained by numerical solution of the boundary value problem given by Eqs. (23)–(27) sharply increase toward the axon terminal in the anterograde-diffusion transport model. This indicates that the anterograde-diffusion transport model is capable of simulating axonal transport of cargo against its concentration gradient. However, in the zeroth approximation, which corresponds to the situation when diffusivity of cargo is negligible, 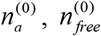, and 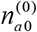 (shown by bold lines in Fig. 7a) remain virtually constant along the axon length. This means that if cargo diffusivity is small, which is the expected situation for slow axonal transport cargos, the anterograde-diffusion transport model is not capable of simulating cargo transport against its concentration gradient.

**Fig. 7.**
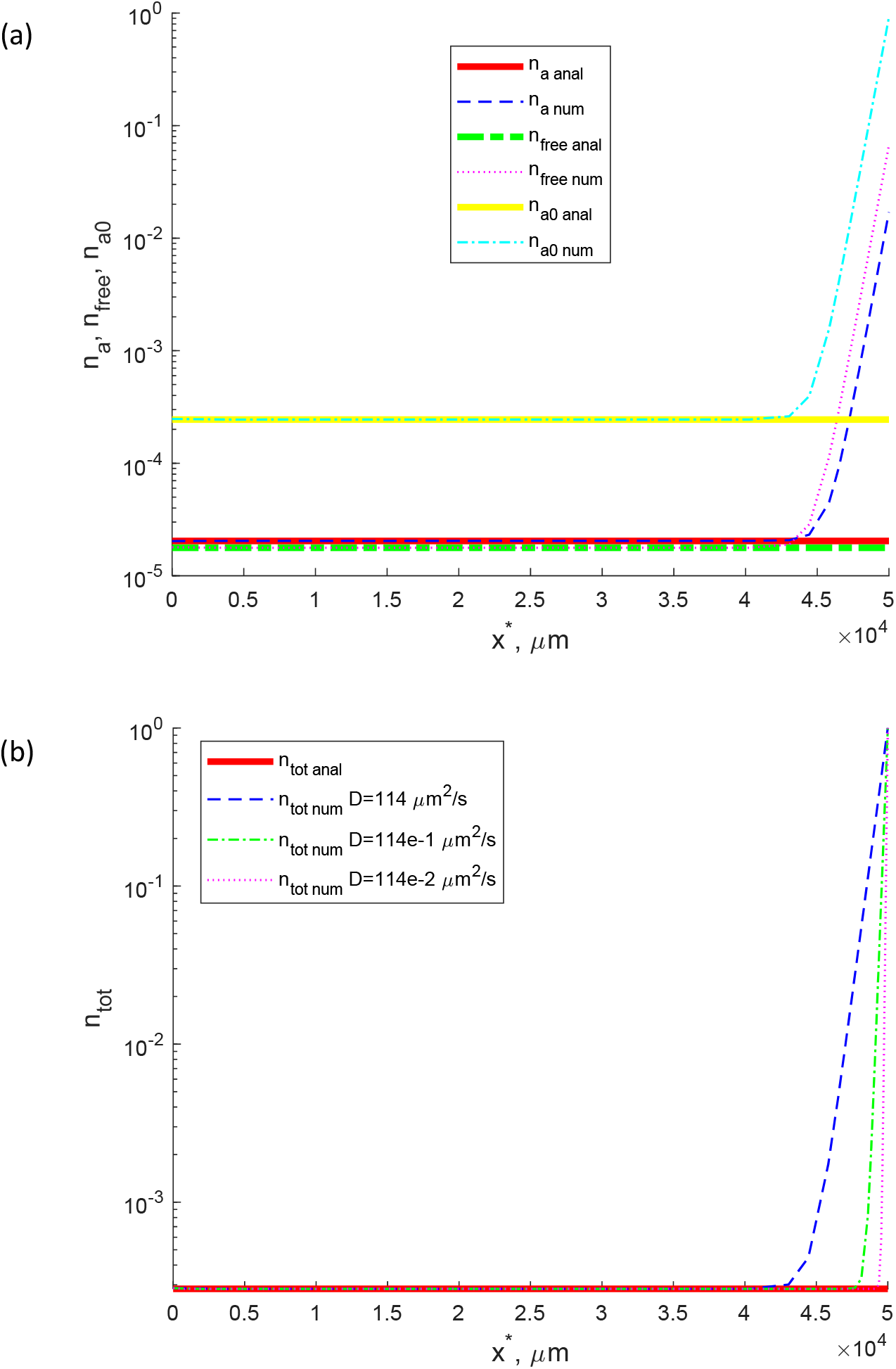
(a) Various components of the total concentration of α-syn monomers. (b) The total concentration of α-syn monomers for the anterograde-diffusion transport model. Bold lines show the analytical solution obtained for the case of small cargo diffusivity. 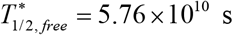 and 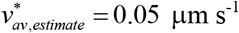.

The total concentration of α-syn monomers (the sum of α-syn monomers concentrations in anterogradely running, free, and pausing kinetic states) can describe α-syn distribution corresponding to cargo transport against the cargo concentration gradient if non-zero diffusivity of α-syn monomers is used. However, if a value of diffusivity is decreased, the diffusion boundary layer near the axon terminal becomes thinner and thinner. In the limit of negligible diffusivity, the concentration is given by a horizontal line (Fig. 7b). This reconfirms our hypothesis that the anterograde-diffusion transport model cannot describe cargo transport against the cargo concentration gradient if cargo diffusivity is very small.

We then tested if a decrease of α-syn half-life to the value expected for α-syn monomers 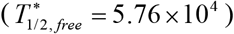 can help in simulating α-syn transport in the axon. The components of the total concentration of α-syn behave similarly to the case with an almost infinite half-life of α-syn. The only difference is a small gradual decay of *n*_*a*0_. The analytical solution corresponding to the case with 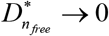 is now a gradually declining curve rather than a horizontal line (compare Fig. 8a and Fig. 7a).

**Fig. 8.**
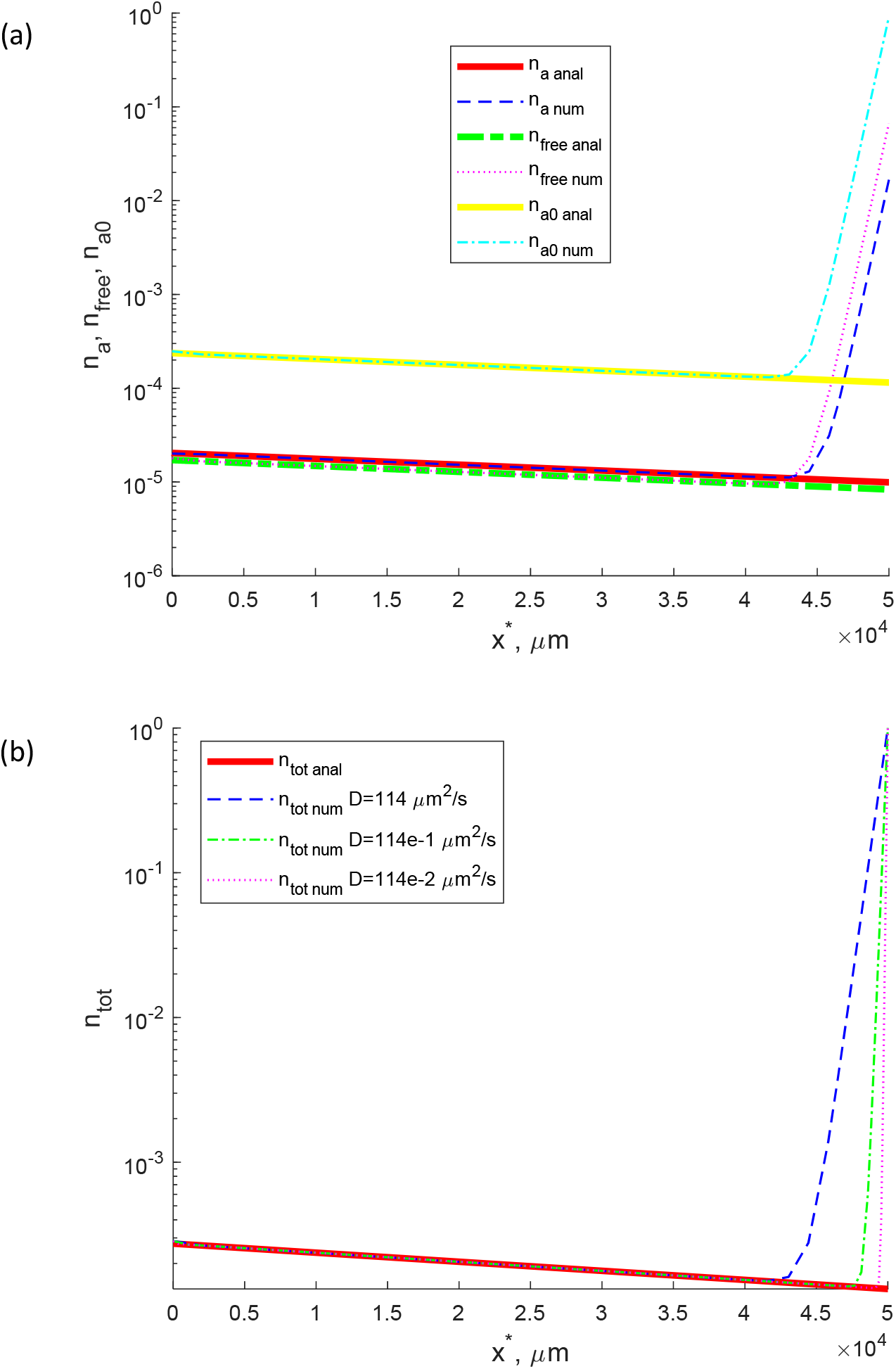
(a) Various components of the total concentration of α-syn monomers for the anterograde-diffusion transport model. (b) The total concentration of α-syn monomers for the anterograde-diffusion transport model. Bold lines show the analytical solution obtained for the case of small cargo diffusivity. 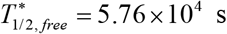 and 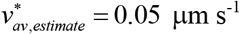.

The total concentration of α-syn monomers for 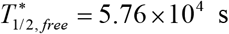 exhibits a small decrease toward the axon terminal (Fig. 8b), which is due to more α-syn destruction during its transport in the axon. The numerical solution for the anterograde-diffusion transport model clearly shows a diffusion boundary layer near the axon terminal. The thickness of this boundary layer decreases with a decrease of α-syn diffusivity. The analytical solution, which describes the limiting case of 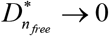 (thick red line in Fig. 8b), shows the inability of the model with anterograde motor-driven transport alone to describe cargo transport against its concentration gradient without the help of diffusion.

### 3.3. The model with anterograde and retrograde motor-driven transport alone (without cargo diffusion) can describe cargo transport against its concentration gradient

If cargo is transported only by anterogradely and retrogradely moving motors, with no free or pausing states (Fig. 2c), the results presented in Figs. S11 and S12 show that the model can still fit the prescribed concentration profile (Fig. S11a) as well as reproduce the prescribed average cargo velocity (Fig. S12a). This is explained by the fact that cargo transport is bidirectional in this case, although cargo diffusion is neglected, and Eqs. (41) and (42) allow for the imposition of given cargo concentrations both at the axon hillock and the axon tip (Eqs. (43) and (44)). The best-fit values of kinetic constants for 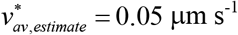 are given in Table S3. These values are constant; no modulation of the kinetic constants is required. The cargo fluxes displayed in Fig. S11b are identical to those displayed in Fig. 3b for the three slow axonal transport cases. This is because the total cargo flux equals the average cargo velocity times the total cargo concentration (Eq. (9)). The anterograde and retrograde cargo fluxes both increase in magnitude at the axon tip (Fig. S12b).

## 4. Discussion, limitations of the model, and future directions

In this paper, we compared bidirectional and unidirectional axonal transport models with respect to whether or not these models allow the imposition of cargo concentration on the tip of the axon. To the best of our knowledge, the results reported in this paper present the first explanation for the utilization of seemingly inefficient bidirectional transport in neurons. The presented analysis uncovers an important feature of bidirectional and unidirectional axonal cargo transport processes. Bidirectional axonal transport includes both anterograde and retrograde components, and hence cargo can be transported in both directions. This allows for the imposition of protein concentration not only at the axon hillock but also at the presynaptic terminal. Since any value of protein concentration can be imposed at the presynaptic terminal, bidirectional axonal transport is capable of transporting protein against its concentration gradient, allowing for the situation in which the protein concentration at the beginning of the axon is low and high at the presynaptic terminal. It should be noted that although cargo diffusion can contribute to retrograde transport in this case, the effect of diffusion is confined to the diffusion boundary layer near the axon terminal. The thickness of the diffusion boundary layer decreases if cargo diffusivity in simulations is decreased. The perturbation analysis for the case of very small cargo diffusivity shows that the anterograde-diffusion model is unable to simulate the increase of cargo concentration toward the axon terminal if cargo diffusivity tends to zero.

We simulated an axon with a length of 50 mm. Since the thickness of the diffusion boundary layer is independent of the axon length, for longer axons the diffusion boundary layer will occupy an even smaller portion of the axon length. Thus, bidirectional axonal transport, which includes both anterograde and retrograde motor-driven components, is needed to transport axonal cargo against its concentration gradient.

The presented results suggest why bidirectional transport, despite its much less efficient utilization of molecular motors and larger energy cost, is used by neurons to transport presynaptic proteins from the soma, where these proteins are synthesized, to the presynaptic terminal. We should note that for cargo to be transported in both anterograde and retrograde directions, it is not enough to have cargos that make bidirectional movements occasionally (for example, when one group of motors overpowers another group of motors attached to the cargo, causing a brief motion in the opposite direction). It is necessary to have separate populations of anterogradely and retrogradely moving cargos, which happens, for example, in slow axonal transport.

Our results show that unidirectional (motor-driven) axonal transport alone cannot describe the increase of protein concentration near the presynaptic terminal. However, if the pausing and diffusion-driven (free) states are removed from the model and only anterograde and retrograde motor-driven states are included in the model, the model is still capable of predicting cargo transport against the concentration gradient. Why then do cargos moved by slow axonal transport spend most of their time pausing if the cargo can be moved with a much larger average velocity without pausing? Our explanation is that relying on fast axonal transport would be much more expensive energetically because cargos would have to be moved continuously in a circulation loop, similar to the transport of dense core vesicles in Drosophila motoneurons (Wong et al. 2012). The utilization of pauses in cargo transport saves energy because it eliminates the need for long-range circulation driven by energy-consuming molecular motors. The circulation may have to be repeated many times for the same cargo.

In our simulations, we used constant values of kinetic constants, which are independent of cargo concentration. This means that a single cargo does not see what the other cargos are doing. This does not change the conclusions from our modeling. It does not matter whether the coefficients in differential equations are constants or functions of dependent variables (concentrations). One needs the ability to impose cargo concentrations at both the axon hillock (*x*\* = 0) and the axon tip (*x*\* = *L*\*) to inform the model of what the cargo concentrations at the boundaries are. If governing equations allow only for the imposition of the concentration at the hillock, the model does not know what concentration it should satisfy at the axon tip.

Finally, a note about the ability of diffusion alone to describe cargo transport against the cargo concentration gradient. Kuznetsov and Kuznetsov (2015) studied a cargo (tau protein) diffusion model with a finite cargo half-life in the free state. Cargo can transition from a free state, where it is moved by diffusion, to a pausing state, where it does not move. Figs. 4b and 4d in Kuznetsov and Kuznetsov (2015) show a perfect curve-fit of the total cargo concentration with experimental data, while Figs. 4a and 4c show how the kinetic constant characterizing the rate of cargo attachment to MTs must be modulated to achieve this fit. The results suggest that although a reaction-diffusion model can describe cargo transport against the concentration gradient, it requires an unphysical increase in a value of a kinetic constant along the axon length. This variation of the kinetic constant value with the axon length heavily depends on how long the axon is (compare Figs. 4a and 4c in Kuznetsov and Kuznetsov 2015).

Future work should include experimental testing of the slow axonal transport model of α-syn monomers transport, which is described in section 2.1, including the calibration of model parameters that we presented. Unlike the model developed in Jung and Brown (2009) for neurofilament transport, the model developed in section 2.1 has not been experimentally tested yet.

Future research should also address models in which kinetic constants depend on cargo concentration gradients, such as the model developed in Ciocanel et al. (2020). In such situations the differential equations become nonlinear.

## Acknowledgment

IAK acknowledges the fellowship support of the Paul and Daisy Soros Fellowship for New Americans and the NIH/National Institute of Mental Health (NIMH) Ruth L. Kirchstein NRSA (F30 MH122076-01). AVK acknowledges the support of the National Science Foundation (award CBET-2042834) and the Alexander von Humboldt Foundation through the Humboldt Research Award.

## Supplemental Materials

### S1. Supplementary tables: best-fit values of kinetic constants for various models

**Table S1.**
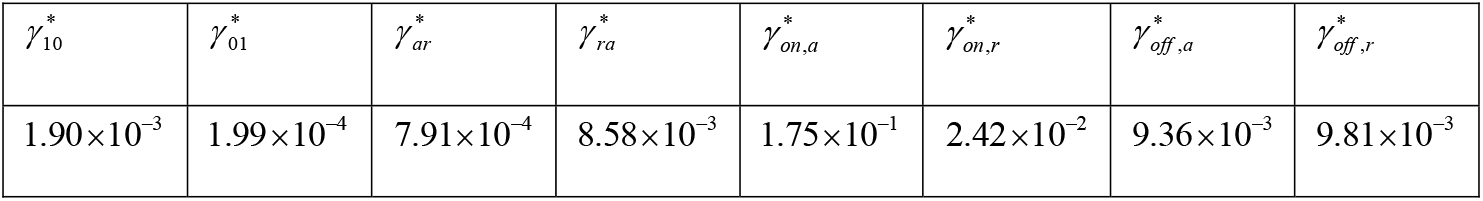
Kinetic constants characterizing the transition of α-syn monomers between different kinetic states in slow axonal transport. 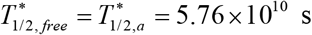 and 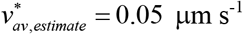 (one of the cases displayed in Figs. 4 and 5, other cases in these figures correspond to 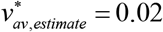 and 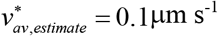). Note that since these kinetic constants are found as best-fit parameters by minimizing the objective function defined by Eq. (55), the values of these constants depend on parameter values, such as 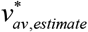. Kinetic constants given in Table S1 were calculated for 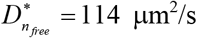. A different set of parameter values would give a different set of kinetic constants. 1000 random points plus the initial point given in Table 3 were utilized as starting points in the search for the global minimum. All kinetic constants in Table S1 have dimensions s^−1^.

**Table S2.** Kinetic constants characterizing the transition of α-syn monomers between different kinetic states in an axonal transport model that includes anterograde motor-driven transport and diffusion only. 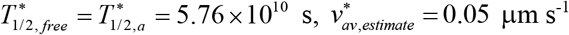, and 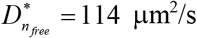. The computational procedure used to obtain these constants was similar to that used for Table S1. All kinetic constants in Table S2 have dimensions s^−1^.

|  |  |  |  |
| --- | --- | --- | --- |
| $\gamma_{10}^*$ | $\gamma_{01}^*$ | $\gamma_{on,a}^*$ | $\gamma_{off,a}^*$ |
| $3.23 \times 10^{-4}$ | $2.69 \times 10^{-5}$ | $1.34 \times 10^{-1}$ | $9.89 \times 10^{-3}$ |

**Table S3.** Kinetic constants characterizing the transition of α-syn monomers between different kinetic states in an axonal transport model that includes anterograde and retrograde motor-driven transport but no diffusion. 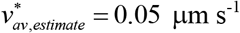. The computational procedure used to obtain these constants was similar to that used for Table S1. All kinetic constants in Table S3 have dimensions s^−1^.

|  |  |
| --- | --- |
| $\gamma_{ar}^*$ | $\gamma_{ra}^*$ |
| $7.02 \times 10^{-3}$ | $8.12 \times 10^{-3}$ |

### S2. Supplementary analysis

#### S2.1. Bidirectional model can describe transport against the concentration gradient for different average velocities of α-syn monomers: additional figures

If α-syn is somehow protected from degradation during its axonal transport and its half-life is very large, the components of the total concentration are constant along the axon length and increase sharply only near the presynaptic terminal (Fig. S1). *n*_*a*0_ is the largest component, followed by *n*_*r*0_, *n_a_*, *n_r_*, and *n_free_*. *n*_*a*0_ and *n*_*r*0_ are the largest because in slow axonal transport the protein spends most of its time in the pausing states (Fig. S1, 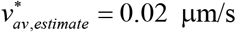). An increase of the average protein velocity results in an increase of the concentration of kinesin-driven protein, *n_a_*, and an increase in the difference between kinesin (anterograde) and dynein (retrograde)-driven protein concentrations, *n_a_* – *n_r_* (Figs. S2 and S3).

**Fig. S1.**
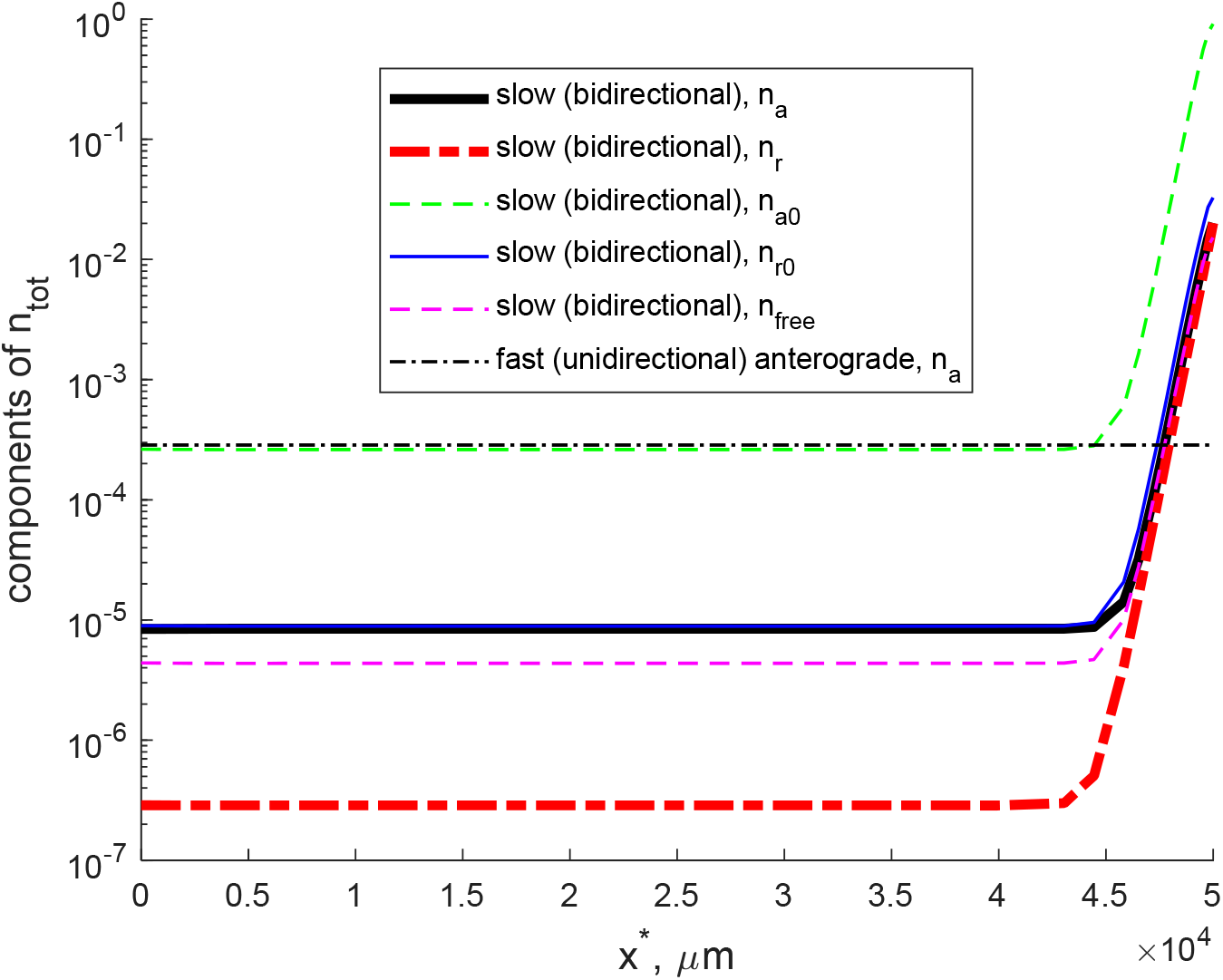
Components of the total concentration of α-syn: *n_a_*, *n_r_*, *n*_*a*0_, *n*_*r*0_, and 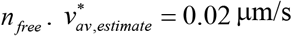 and 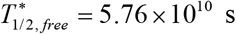. The fraction of α-syn monomers conveyed in the fast component of axonal transport is assumed to be zero.

**Fig. S2.**
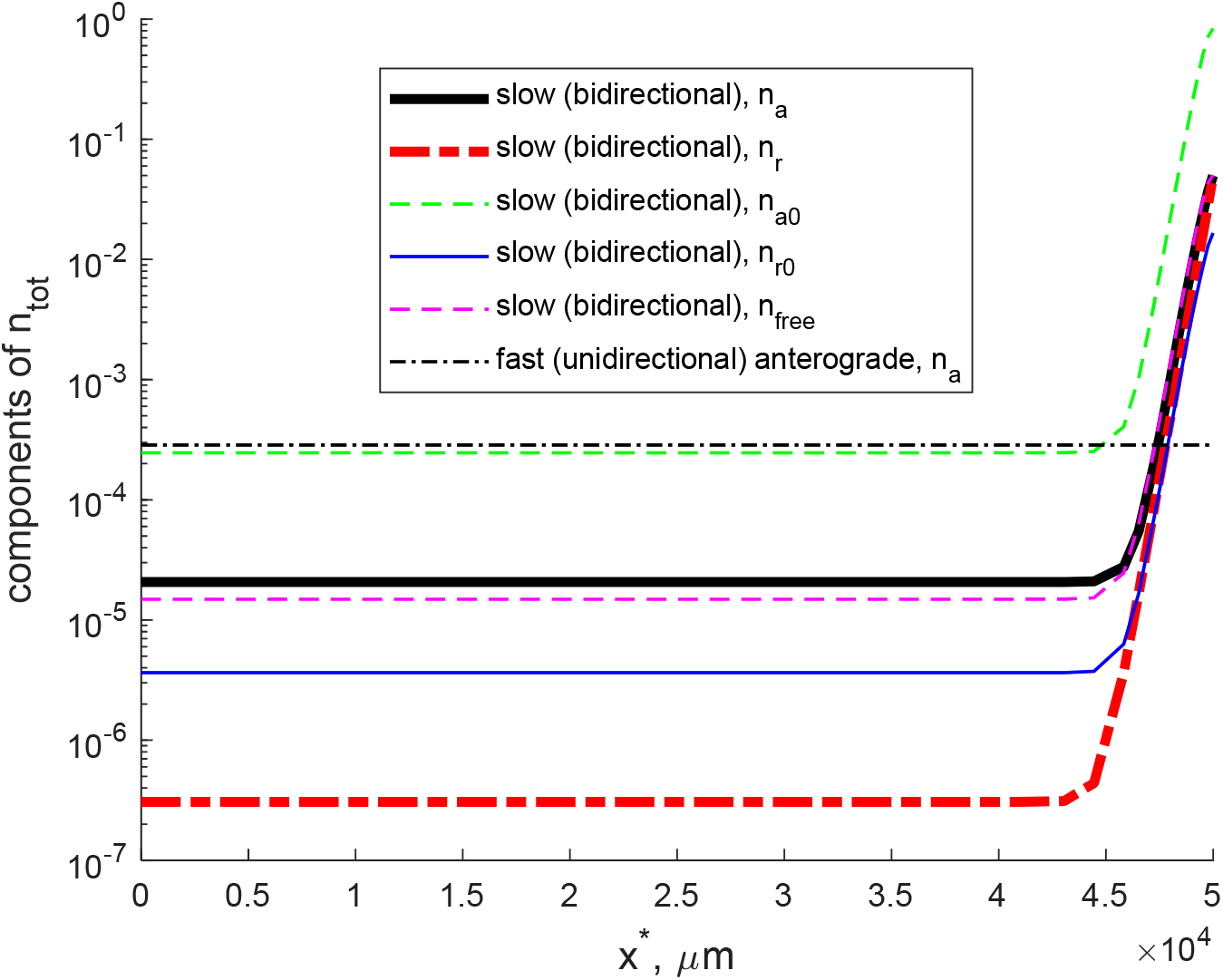
Components of the total concentration of α-syn: *n_a_*, *n_r_*, *n*_*a*0_, *n*_*r*0_, and 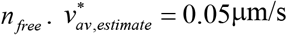 and 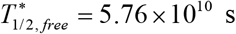. The fraction of α-syn monomers conveyed in the fast component of axonal transport is assumed to be zero.

**Fig. S3.**
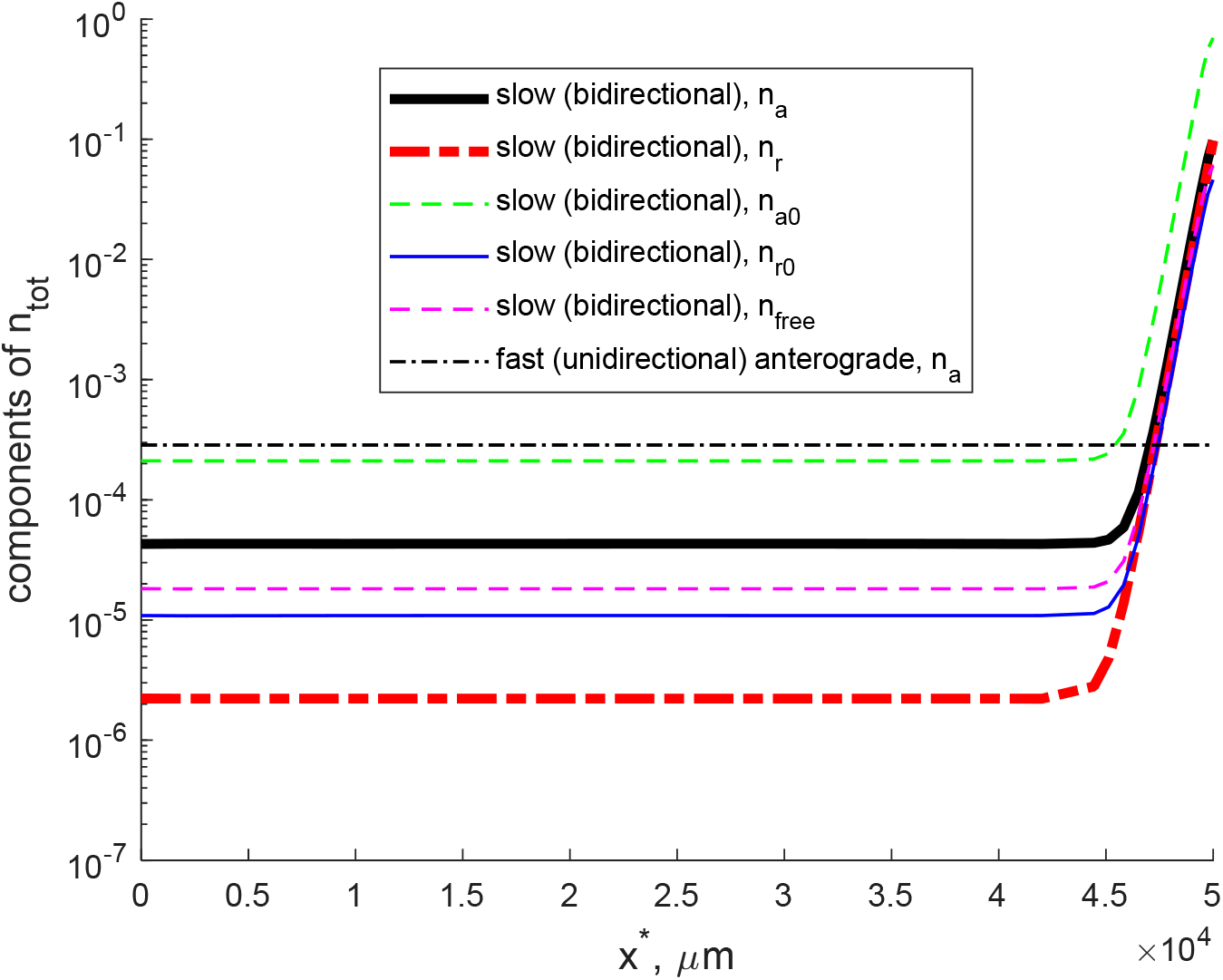
(a) Components of the total concentration of α-syn: *n_a_*, *n_r_*, *n*_*a*0_, *n*_*r*0_, and 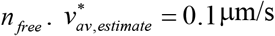 and 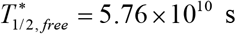. The fraction of α-syn monomers conveyed in the fast component of axonal transport is assumed to be zero.

#### S2.2. Bidirectional model can describe transport against the concentration gradient for different half-lives of α-syn monomers

If the α-syn half-life is varied, the slow axonal transport model can still fit an α-syn concentration profile with a sharp concentration increase at the presynaptic terminal (the total concentration is shown in Fig. S4a, the components of the total concentration are shown in Figs. S5–S7). The fast axonal transport model predicts a decreasing concentration toward the presynaptic terminal due to protein degradation. The decrease becomes faster for a shorter protein half-life (Fig. S5b vs Figs. S6b and S7b).

For the slow axonal transport model, the α-syn flux remains almost constant along most of the axon (Fig. 7b). This is because in the slow axonal transport model we assumed that α-syn is destroyed only in the free (cytosolic) kinetic state (see Eq. (5)), and the concentration of α-syn in this kinetic state is very low in most of the axon (Fig. S5, S6, S7). Close to the presynaptic terminal the flux decreases, which is due to a sharp increase in the concentration of α-syn (including in the free state, in which α-syn can be destroyed) near the presynaptic terminal (Fig. S4a and Figs. S5, S6, S7). For the fast anterograde axonal transport model, α-syn degradation during its transport along the axon results in a decrease of the α-syn flux because of α-syn destruction (Fig. S4b), see Eq. (14).

It is interesting that even if the α-syn half-life is decreased, the slow axonal transport model is capable of simulating a constant velocity along most of the axon, with the velocity decreasing close to the presynaptic terminal (Fig. S8a). The diffusion component of the total flux is much smaller than the anterograde and retrograde components (Fig. S8b).

**Fig. S4.**
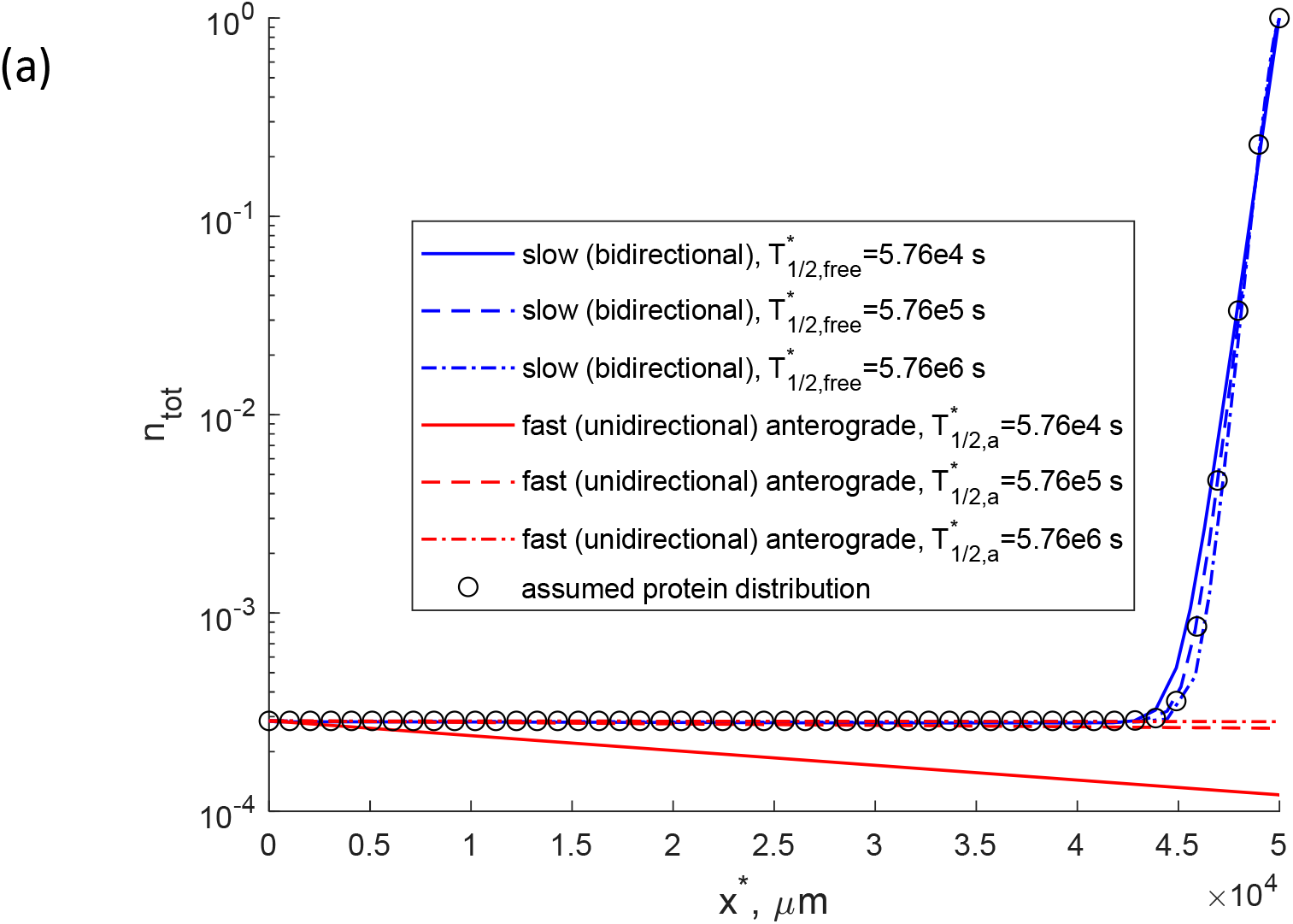
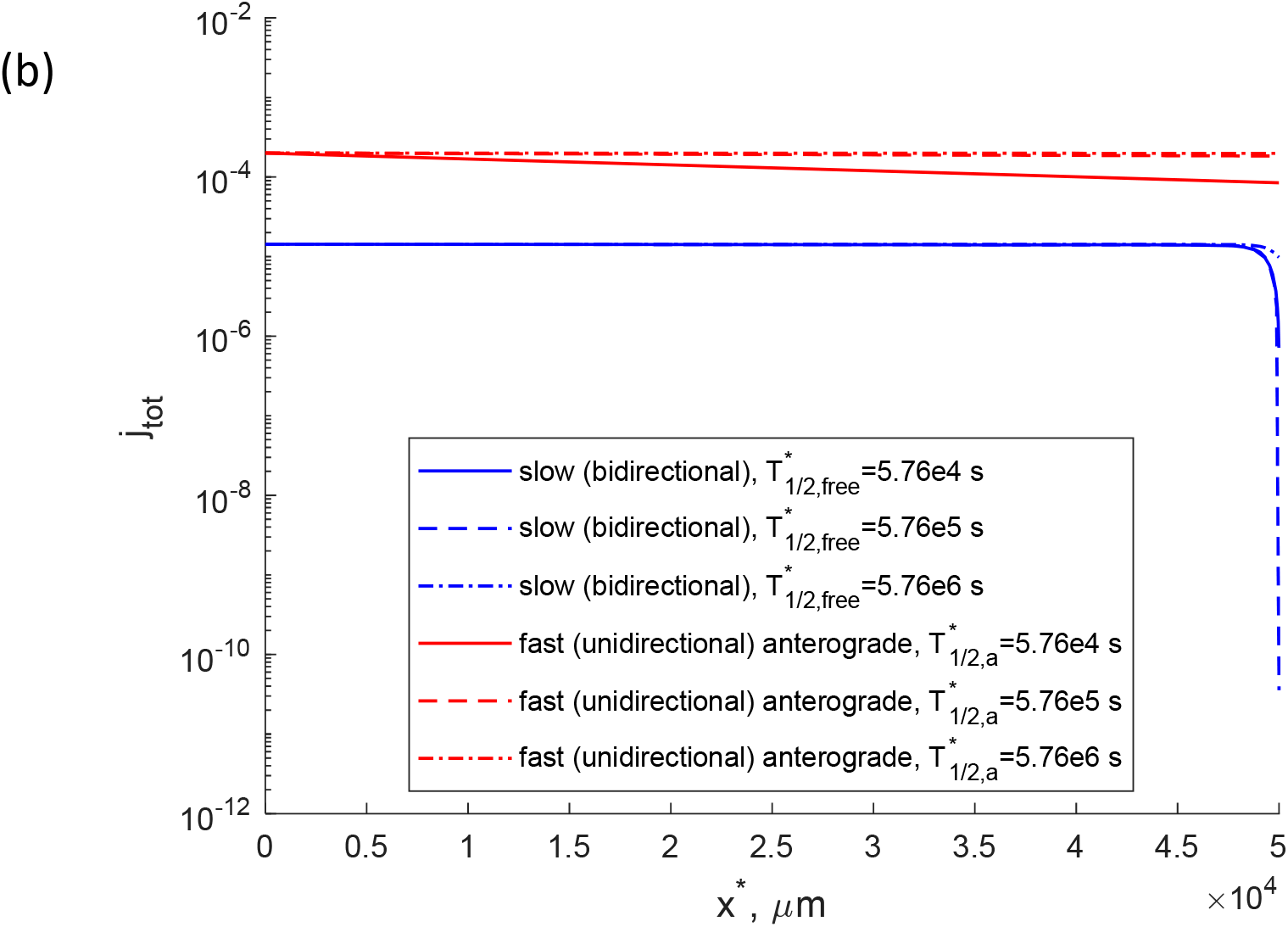
Slow (bidirectional) and fast (unidirectional) axonal transport. 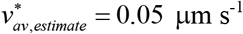. The fraction of α-syn monomers conveyed in the fast component of axonal transport is assumed to be zero. (a) Total concentration of α-syn monomers (the sum of α-syn concentrations in motor-driven, pausing, and diffusing states). (b) Total flux of α-syn monomers due to the action of molecular motors and diffusion in the cytosol.

**Fig. S5.**
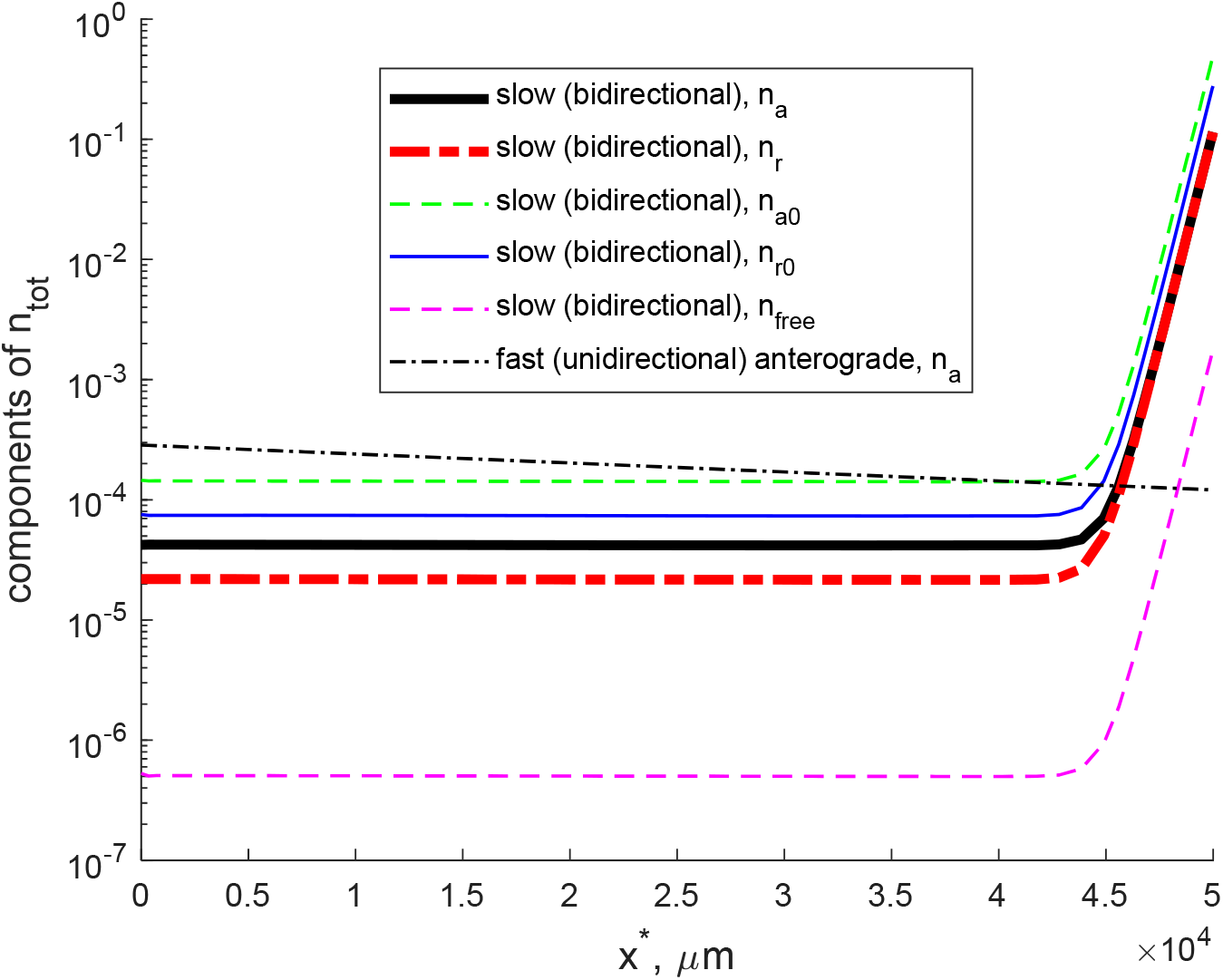
Components of the total concentration of α-syn: *n_a_*, *n_r_*, *n*_*a*0_, *n*_*r*0_, and 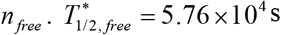 and 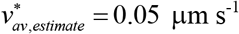. The fraction of α-syn monomers conveyed in the fast component of axonal transport is assumed to be zero.

**Fig. S6.**
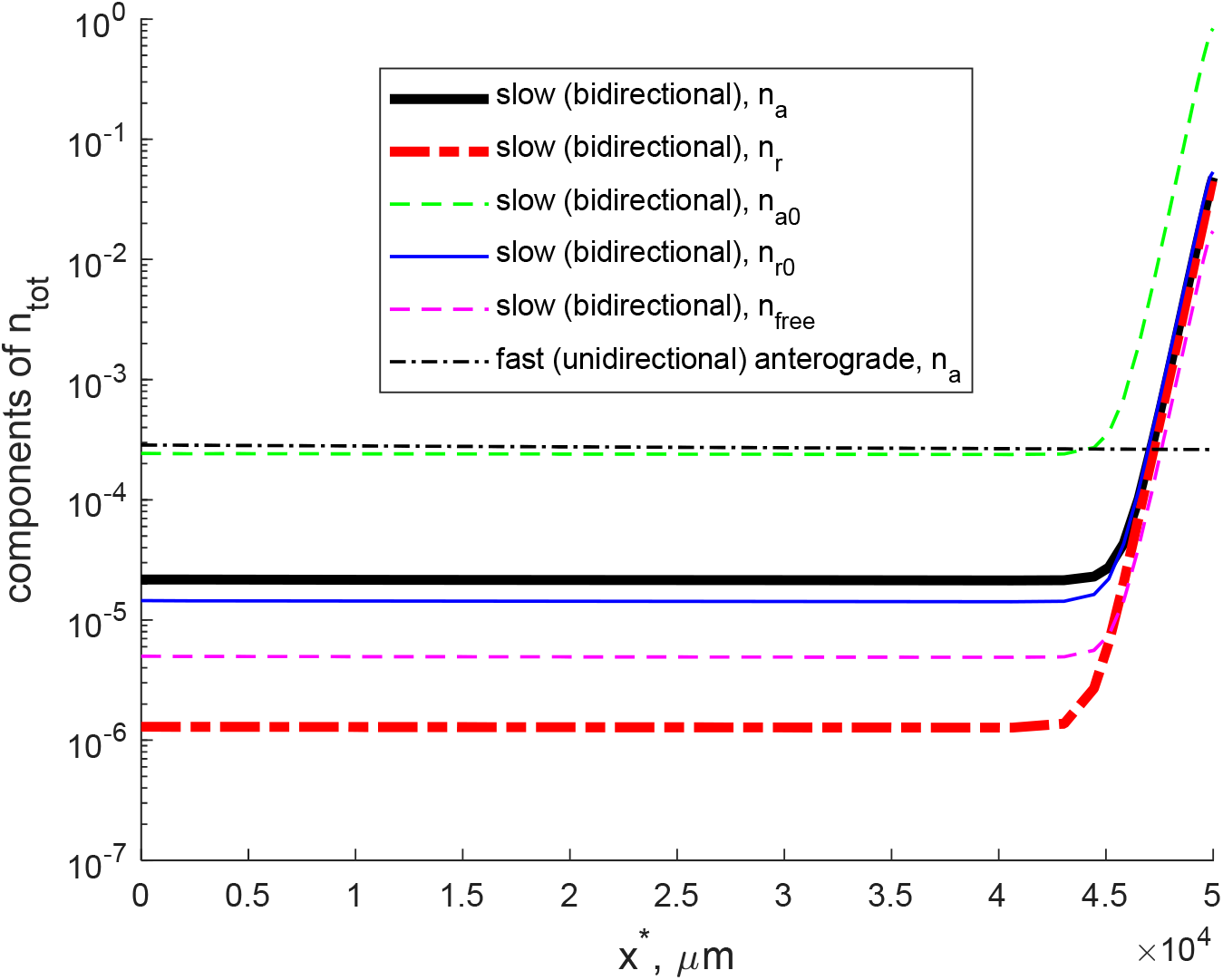
Components of the total concentration of α-syn: *n_a_*, *n_r_*, *n*_*a*0_, *n*_*r*0_, and 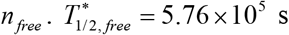 and 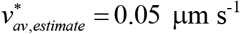. The fraction of α-syn monomers conveyed in the fast component of axonal transport is assumed to be zero.

**Fig. S7.**
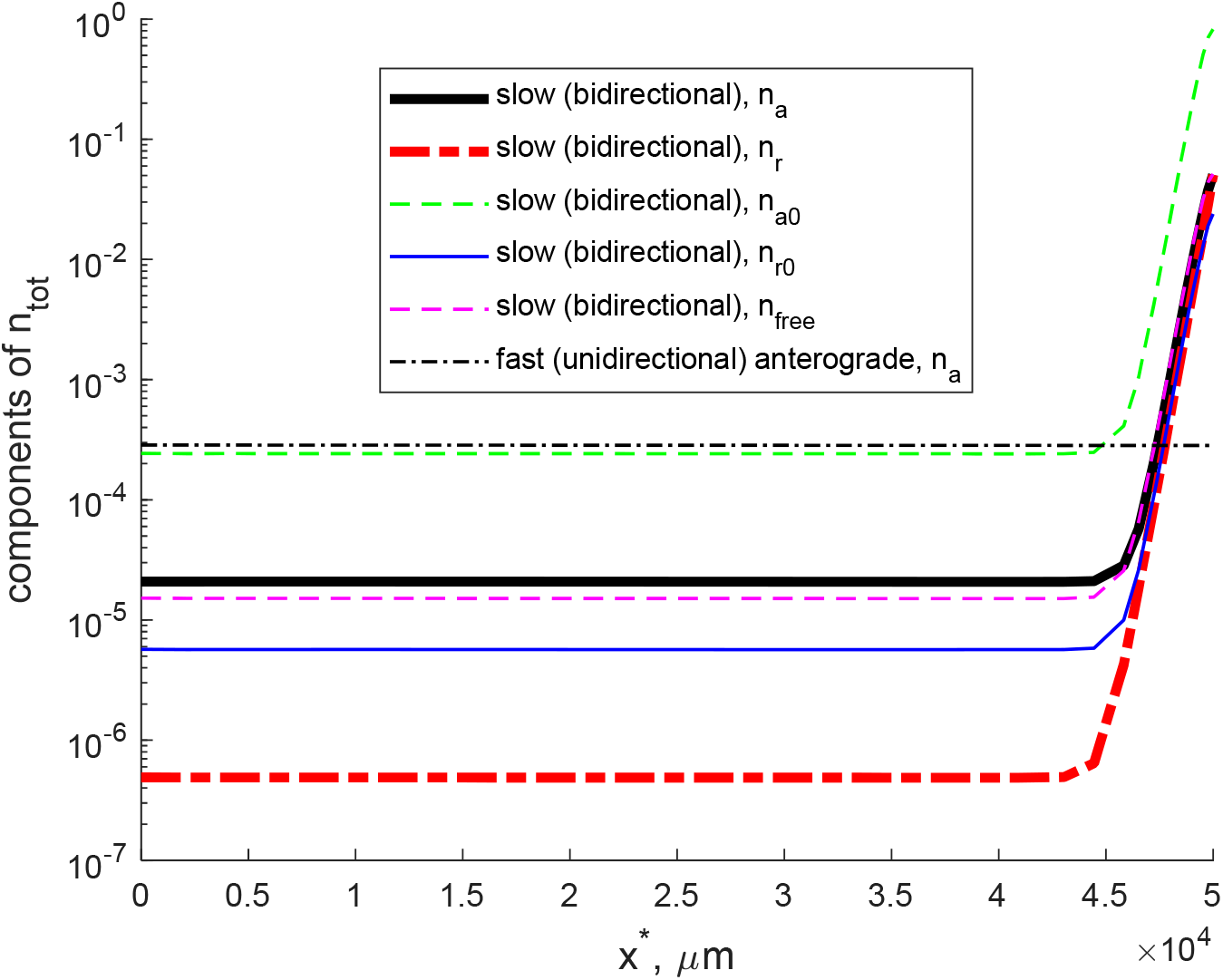
Components of the total concentration of α-syn: *n_a_*, *n_r_*, *n*_*a*0_, *n*_*r*0_, and 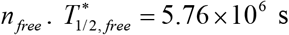 and 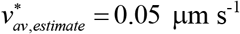. The fraction of α-syn monomers conveyed in the fast component of axonal transport is assumed to be zero.

**Fig. S8.**
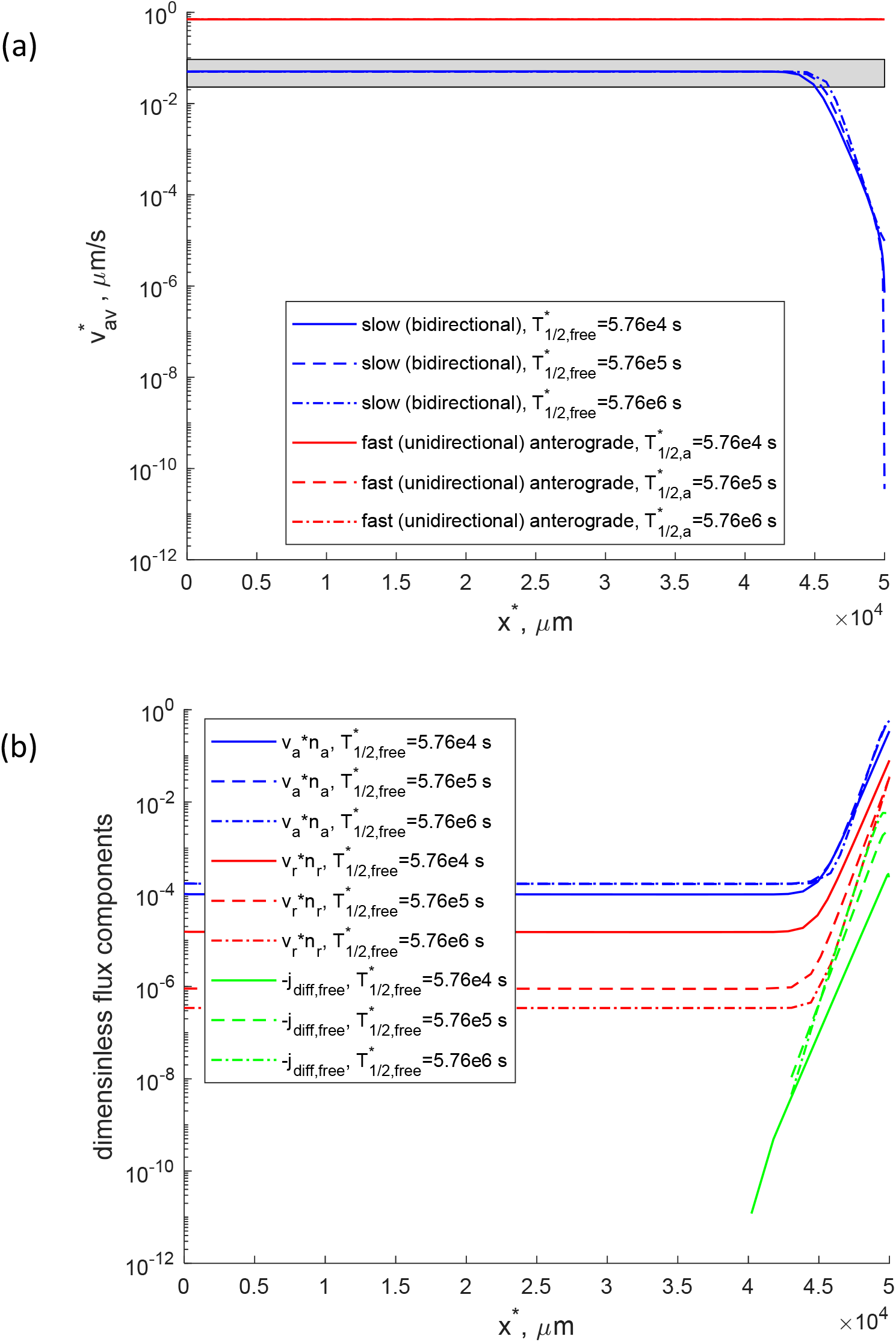
Slow (bidirectional) and fast (unidirectional) axonal transport. 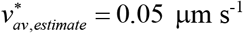. The fraction of α-syn monomers conveyed in the fast component of axonal transport is assumed to be zero. (a) Average velocity of α-syn monomers in slow axonal transport along the axon. (b) Anterograde motor-driven, retrograde motor-driven, and diffusion-driven components of the total α-syn flux. Note that because of the log scale used, Fig. S8b shows the absolute values of the flux components. Since retrograde and diffusion-driven flux components are negative, Fig. 8b shows the negative of these components.

#### S2.3. Bidirectional model can describe transport against the concentration gradient for the situation when a 15% fraction of α-syn monomers is moved by fast axonal transport

To investigate whether transport of a fraction of α-syn by fast axonal transport may affect the α-syn distribution, we combined the slow and fast components of axonal transport and modified the boundary conditions in such a way that 85% of α-syn enters the axon in the slow component and 15% enters the axon by axonal transport (Tang et al. 2012; Roy 2014), see Eqs. (21) and (22). The combined 15% fast plus 85% slow axonal transport model is still able to fit the α-syn concentration characterized by a uniform concentration in most of the axon, with a concentration increase only at the presynaptic terminal (Fig. S9a).

The α-syn flux in the case of combined fast plus slow axon transport is slightly (by ~17%) larger because fast axonal transport is much more efficient than slow axonal transport; even 15% makes a difference (Fig. S9b). The average velocity of α-syn transport is also slightly (by ~15%) larger when a fraction of α-syn is transported in the fast component (Fig. S10a). Cargo transport is governed by the balance of anterograde and retrograde motor-driven transport, with diffusion-driven transport being negligible (Fig. S10b).

**Fig. S9.**
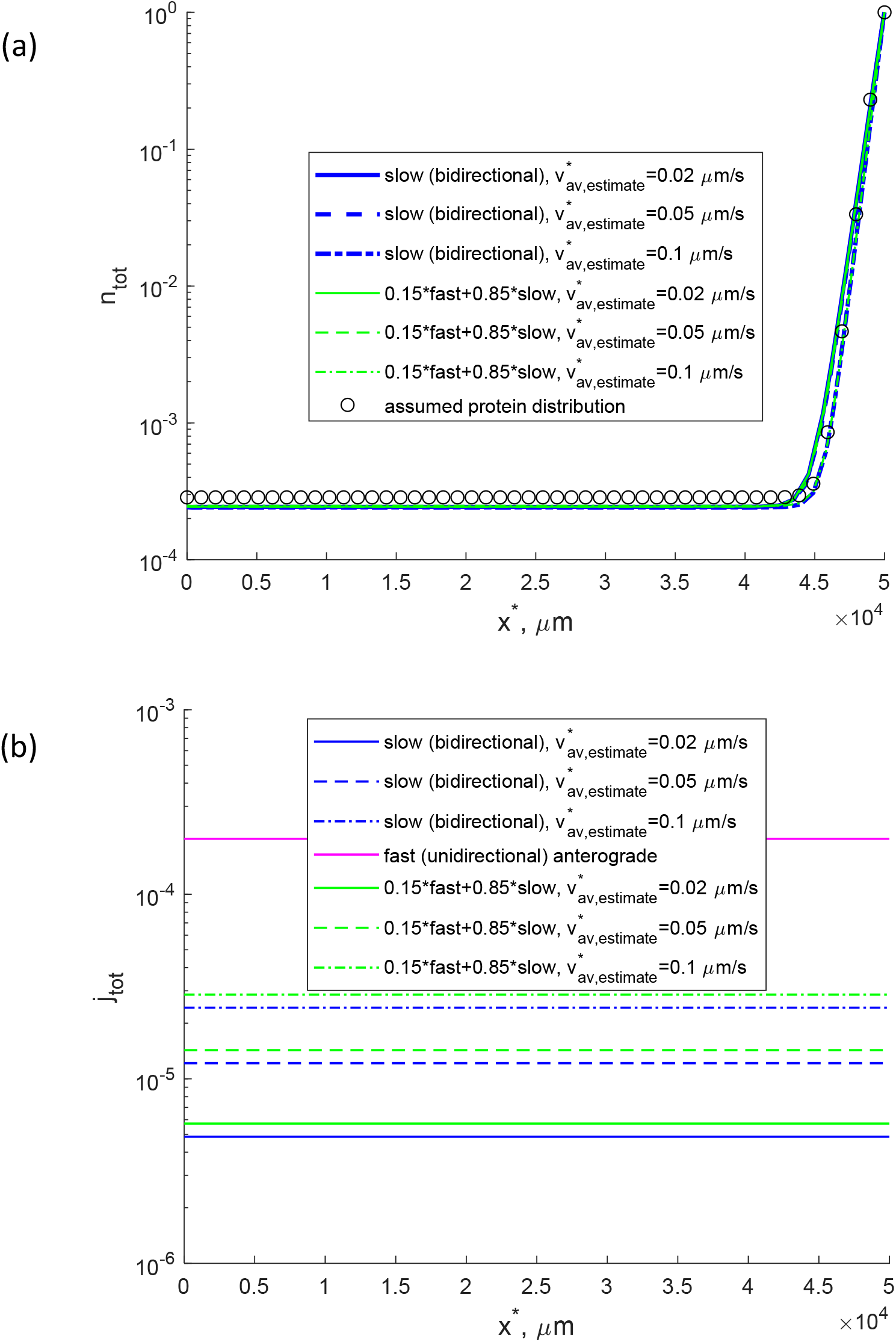
Slow (bidirectional) and fast (unidirectional) axonal transport. 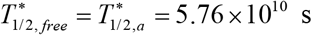 and 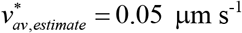. The fraction of α-syn monomers conveyed in the fast component of axonal transport is assumed to be 15%. (a) Total concentration of α-syn monomers (the sum of α-syn concentrations in motor-driven, pausing, and diffusing states). (b) Total flux of α-syn monomers due to the action of molecular motors and diffusion in the cytosol.

**Fig. S10.**
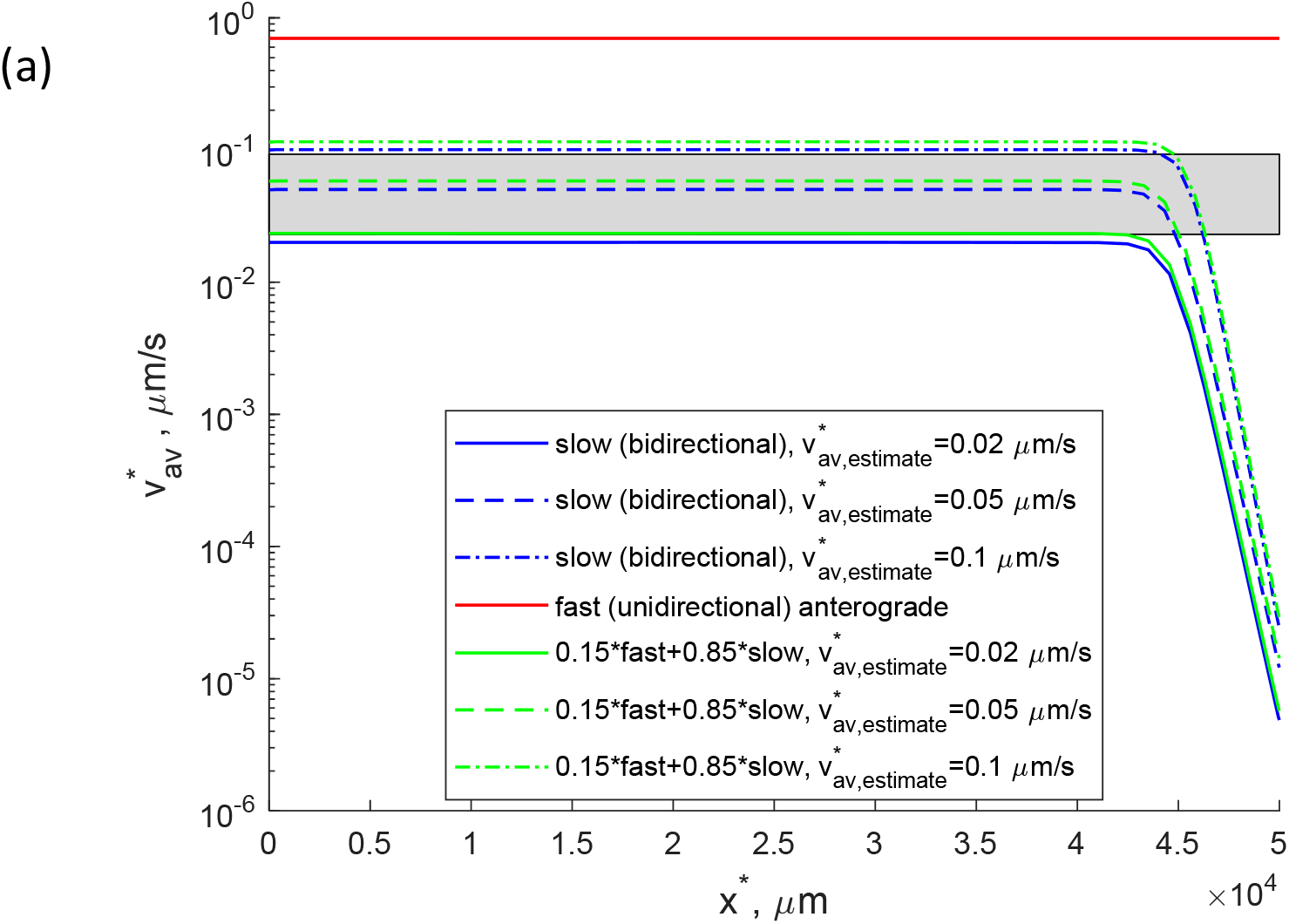
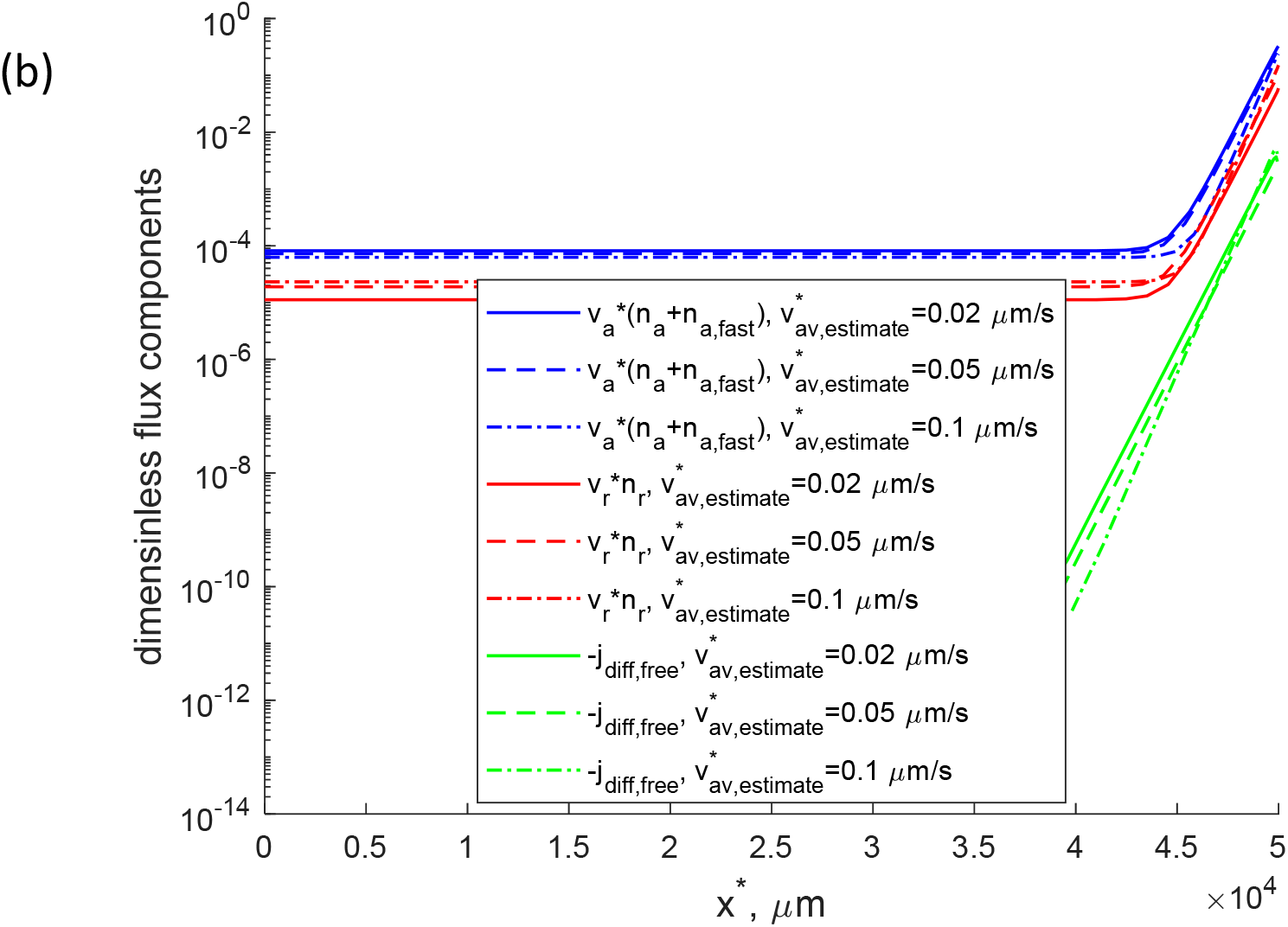
Slow (bidirectional) and fast (unidirectional) axonal transport. 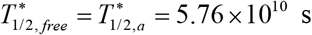 and 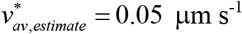. The fraction of α-syn monomers conveyed in the fast component of axonal transport is assumed to be 15%. (a) Average velocity of α-syn monomers in slow (bidirectional) and fast (unidirectional) axonal transport along the axon. (b) Anterograde motor-driven, retrograde motor-driven, and diffusion-driven components of the total α-syn flux. *n_a_* and *n_a, fast_* are scaled in such a way so that 85% of α-syn monomers are transported in the slow component and 15% are transported in the fast component. Note that because of the log scale used, Fig. S10b shows the absolute values of the flux components. Since retrograde and diffusion-driven flux components are negative, Fig. 10b shows the negative of these components.

#### S2.4. Bidirectional model can describe transport against the concentration gradient if cargo diffusivity is removed from the model and only motor-driven transport is retained

Figs. S11 and S12 show that including the diffusivity of cargo is not required to describe axonal transport against the concentration gradient. A model that accounts for motor-driven transport in both directions but does not include cargo diffusion can describe cargo transport toward the axon tip with a higher cargo concentration at the tip of the axon than at the hillock.

**Fig. S11.**
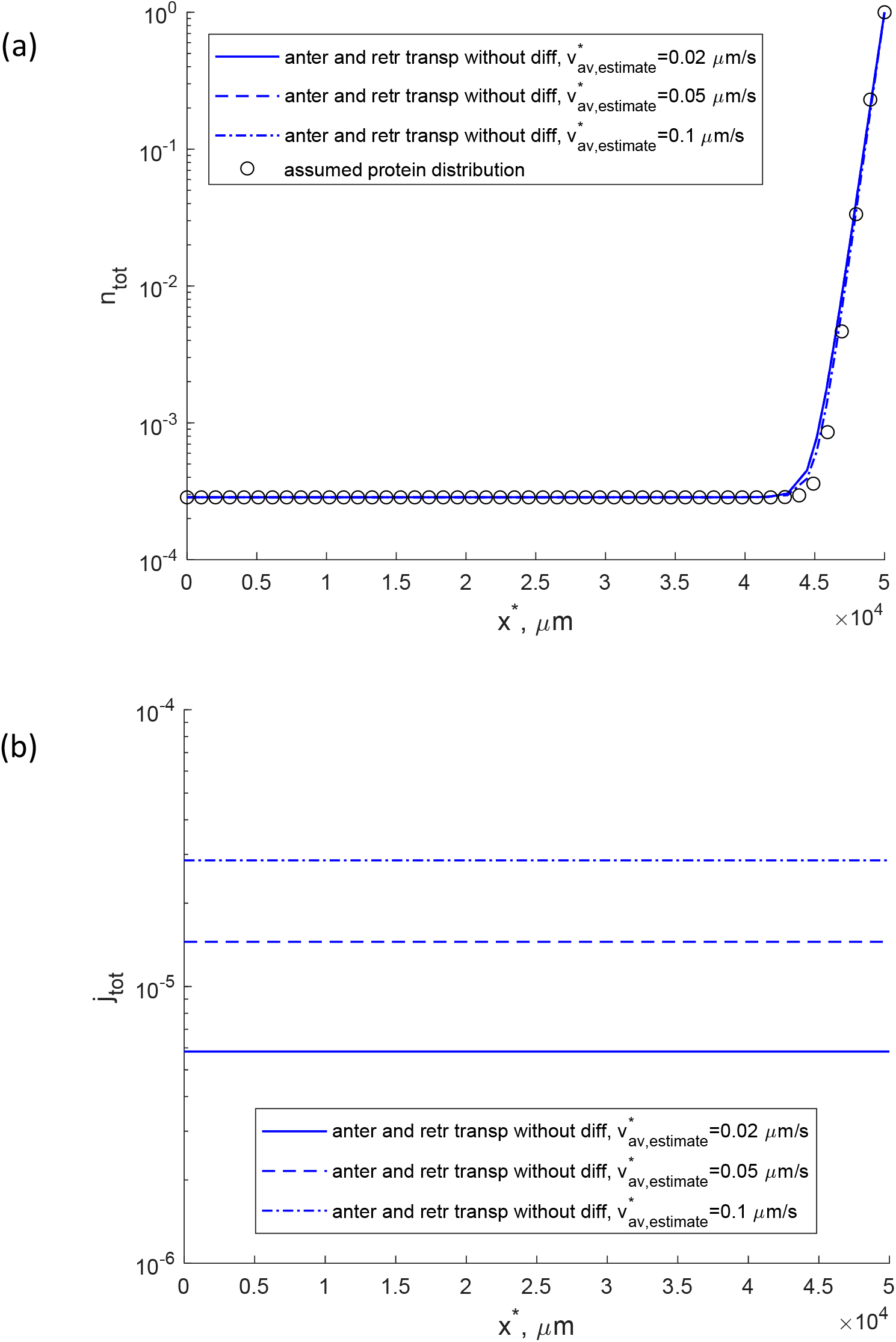
Anterograde and retrograde axonal transport model without pausing states and diffusion. (a) Total cargo concentration (the sum of cargo concentrations in anterograde motor-driven, pausing, and diffusing states). (b) Total flux of cargo due to the action of anterograde and retrograde motors.

**Fig. S12.**
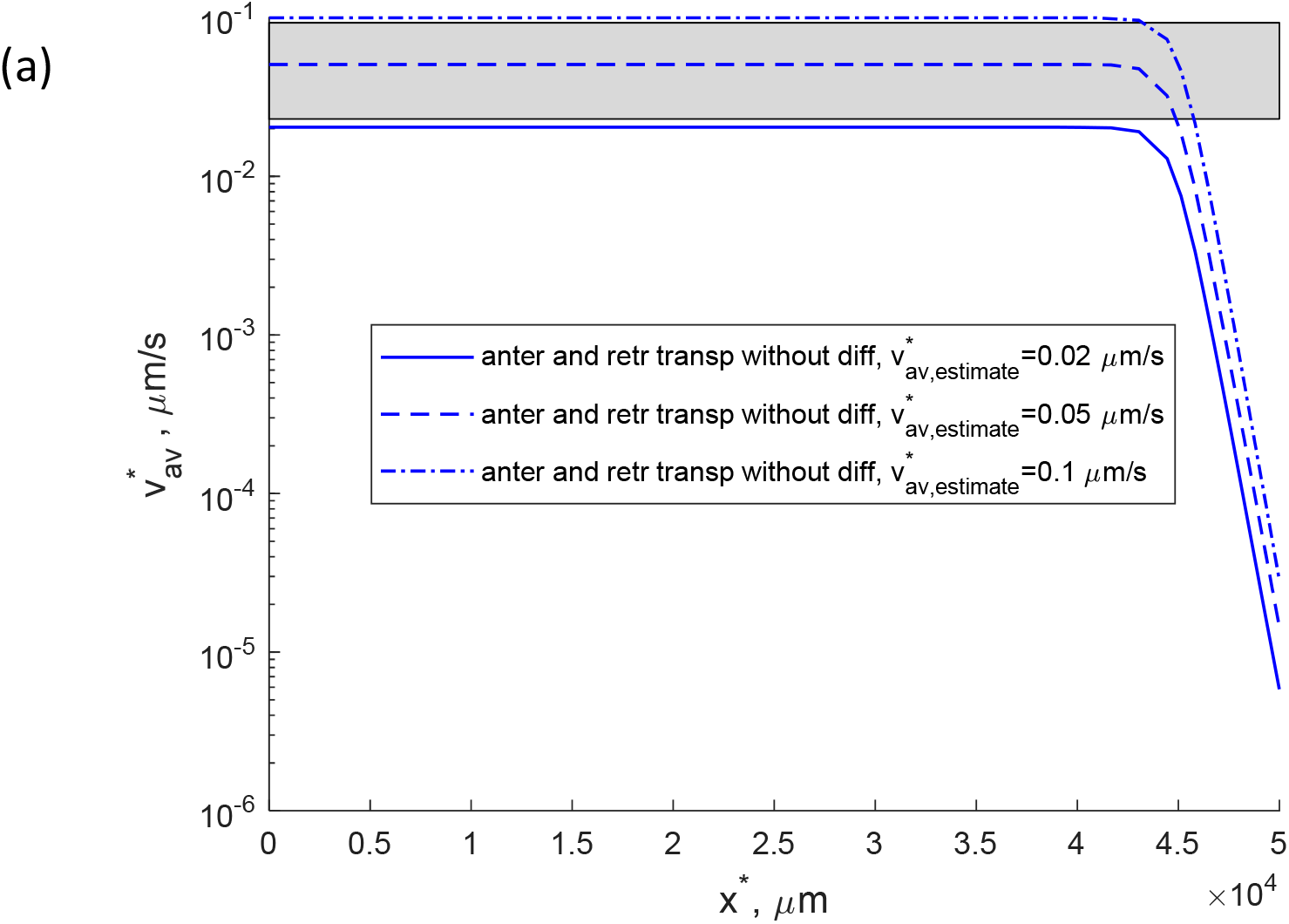
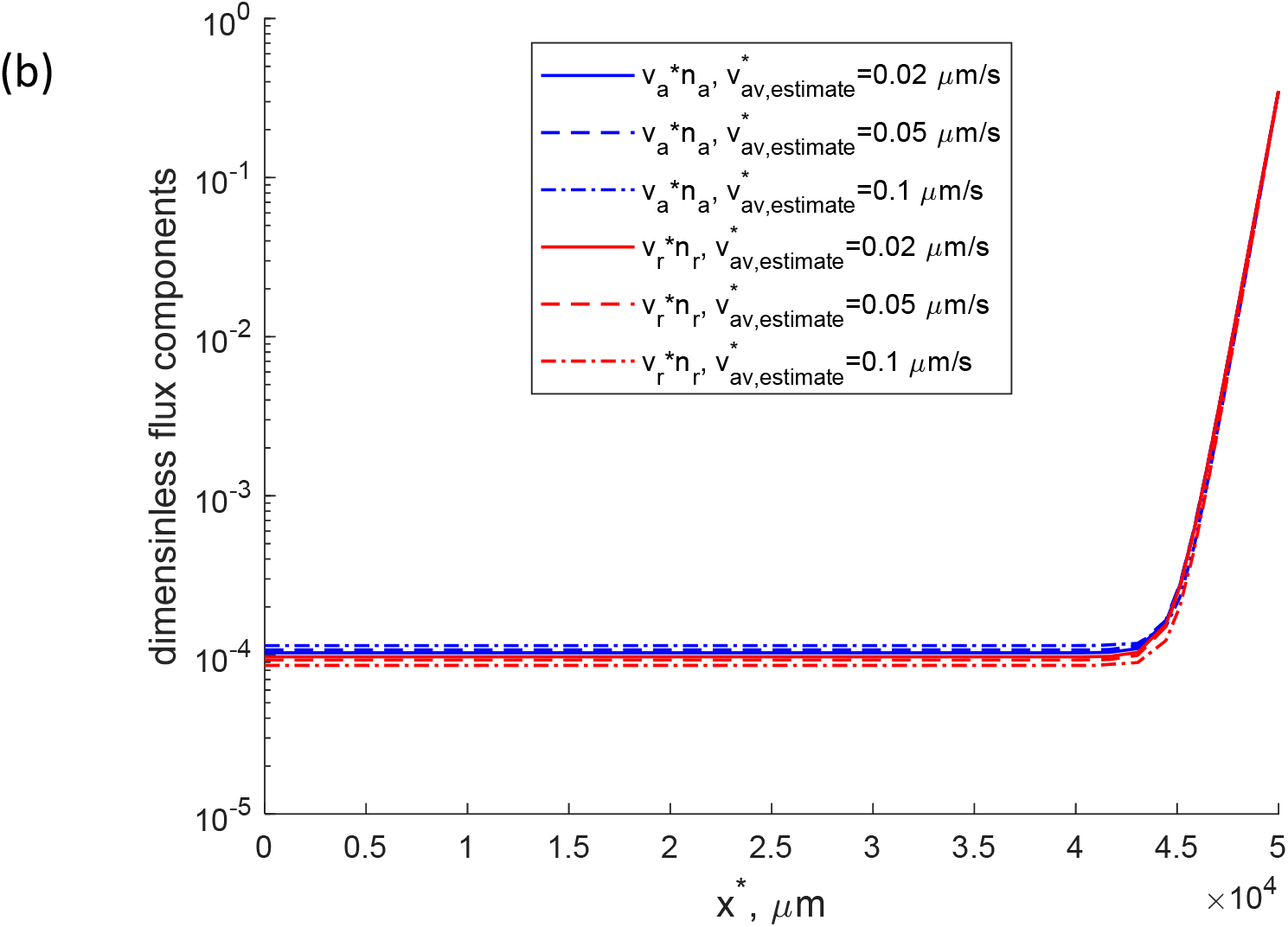
Anterograde and retrograde axonal transport model without pausing states and diffusion. (a) Average cargo velocity. (b) Anterograde motor-driven and retrograde motor-driven components of the total α-syn flux. Note that because of the log scale used, Fig. S12b shows the absolute values of the flux components. Since the retrograde flux component is negative, Fig. 12b shows the negative of this component.

## References

Banks, S.M.L., Medeiros, A.T., McQuillan, M., Busch, D.J., Ibarraran-Viniegra, A.S., Sousa, R., Lafer, E.M., Morgan, J.R., 2020. Hsc70 ameliorates the vesicle recycling defects caused by excess alpha-synuclein at synapses. Eneuro 7, 0448–19.2020.

Beck, J.V. and Arnold, K.J., 1977. Parameter Estimation in Science and Engineering, Wiley, New York.

Bennett, M.C., Bishop, J.F., Leng, Y., Chock, P.B., Chase, T.N., Mouradian, M.M., 1999. Degradation of alpha-synuclein by proteasome. Journal of Biological Chemistry 274, 33855–33858.

Black, M.M., Slaughter, T., Moshiach, S., Obrocka, M., Fischer, I., 1996. Tau is enriched on dynamic microtubules in the distal region of growing axons. Journal of Neuroscience 16, 3601–3619.

Bressloff, P.C., Karamched, B.R., 2016. Model of reversible vesicular transport with exclusion. Journal of Physics A-Mathematical and Theoretical 49, 345602.

Brown, A. (2016). Axonal transport. In D. Pfaff, & N. Volkow (Eds.), Neuroscience in the 21st century (pp. 379) Springer. doi:10.1007/978-1-4939-3474-4_14.

Burre, J., 2015. The synaptic function of alpha-synuclein. Journal of Parkinsons Disease 5, 699–713.

Charvin, D., Medori, R., Hauser, R.A., Rascol, O., 2018. Therapeutic strategies for Parkinson disease: Beyond dopaminergic drugs. Nature Reviews Drug Discovery 17, 804–822.

Ciocanel, M., Jung, P., Brown, A., 2020. A mechanism for neurofilament transport acceleration through nodes of ranvier. Molecular Biology of the Cell 31, 640–654.

Fortin, D., Nemani, V., Voglmaier, S., Anthony, M., Ryan, T., Edwards, R., 2005. Neural activity controls the synaptic accumulation of alpha-synuclein. Journal of Neuroscience 25, 10913–10921.

George, J.L., Mok, S., Moses, D., Wilkins, S., Bush, A.I., Cherny, R.A., Finkelstein, D.I., 2009. Targeting the progression of Parkinson’s disease. Current Neuropharmacology 7, 9–36.

Goldberg, A., 2003. Protein degradation and protection against misfolded or damaged proteins. Nature 426, 895–899.

Gupta, A., Dawson, T.M., 2010. Chapter 10 - pathogenesis of Parkinson’s disease. Blue Books of Neurology 34, 155–169.

Hancock, W.O., 2014. Bidirectional cargo transport: Moving beyond tug of war. Nature Reviews Molecular Cell Biology 15, 615–628.

Hannestad, J.K., Rocha, S., Agnarsson, B., Zhdanov, V.P., Wittung-Stafshede, P., Hook, F., 2020. Single-vesicle imaging reveals lipid-selective and stepwise membrane disruption by monomeric alpha-synuclein. Proceedings of the National Academy of Sciences of the United States of America 117, 14178–14186.

Iyer, A., Claessens, M.M.A.E., 2019. Disruptive membrane interactions of alpha-synuclein aggregates. Biochimica Et Biophysica Acta-Proteins and Proteomics 1867, 468–482.

Jensen, P., Li, J., Dahlstrom, A., Dotti, C., 1999. Axonal transport of synucleins is mediated by all rate components. European Journal of Neuroscience 11, 3369–3376.

Jensen, P., Nielsen, M., Jakes, R., Dotti, G., Goedert, M., 1998. Binding of alpha-synuclein to brain vesicles is abolished by familial Parkinson’s disease mutation. Journal of Biological Chemistry 273, 26292–26294.

Jung, P., Brown, A., 2009. Modeling the slowing of neurofilament transport along the mouse sciatic nerve. Physical Biology 6, 046002.

Karamched, B.R., Bressloff, P.C., 2017. Effects of cell geometry on reversible vesicular transport. Journal of Physics A-Mathematical and Theoretical 50, 055601.

Kim, C., Lee, S., 2008. Controlling the mass action of alpha-synuclein in Parkinson’s disease. Journal of Neurochemistry 107, 303–316.

Kool, J.B., Parker, J.C., van Genuchten, M.T., 1987. Parameter estimation for unsaturated flow and transport models - A review. Journal of Hydrology 91, 255–293.

Kuznetsov, I.A., Kuznetsov, A.V., 2018a. How the formation of amyloid plaques and neurofibrillary tangles may be related – A mathematical modelling study. Proceedings of the Royal Society A 474, 20170777.

Kuznetsov, A.V., Avramenko, A.A., Blinov, D.G., 2009a. Effect of protein degradation in the axon on the speed of the bell-shaped concentration wave in slow axonal transport. International Communications in Heat and Mass Transfer 36, 641–645.

Kuznetsov, A.V., Avramenko, A.A., Blinov, D.G., 2009b. Macroscopic modeling of slow axonal transport of rapidly diffusible soluble proteins. International Communications in Heat and Mass Transfer 36, 293–296.

Kuznetsov, I.A., Kuznetsov, A.V., 2015. A comparison between the diffusion-reaction and slow axonal transport models for predicting tau distribution along an axon. Mathematical Medicine and Biology 32, 263–283.

Kuznetsov, I.A., Kuznetsov, A.V., 2017a. Simulating tubulin-associated unit transport in an axon: Using bootstrapping for estimating confidence intervals of best fit parameter values obtained from indirect experimental data. Proceedings of the Royal Society A 473, 20170045.

Kuznetsov, I.A., Kuznetsov, A.V., 2017b. Utilization of the bootstrap method for determining confidence intervals of parameters for a model of MAP1B protein transport in axons. Journal of Theoretical Biology 419, 350–361.

Kuznetsov, I.A., Kuznetsov, A.V., 2018b. Simulating the effect of formation of amyloid plaques on aggregation of tau protein. Proceedings of the Royal Society A-Mathematical Physical and Engineering Sciences 474, 20180511.

Kuznetsov, I.A., Kuznetsov, A., V., 2020. Modeling tau transport in the axon initial segment. Mathematical Biosciences 329, 108468.

Lashuel, H.A., Overk, C.R., Oueslati, A., Masliah, E., 2013. The many faces of alpha-synuclein: From structure and toxicity to therapeutic target. Nature Reviews Neuroscience 14, 38–48.

Lee, R.H., Mitchell, C.S., 2015. Axonal transport cargo motor count versus average transport velocity: Is fast versus slow transport really single versus multiple motor transport? Journal of Theoretical Biology 370, 39–44.

Li, J.Y., Jensen, P.H., Dahlstrom, A., 2002. Differential localization of alpha-, beta-and gamma-synucleins in the rat CNS. Neuroscience 113, 463–478.

Li, W.X., Hoffman, P.N., Stirling, W., Price, D.L., Lee, M.K., 2004. Axonal transport of human alpha-synuclein slows with aging but is not affected by familial Parkinson’s disease-linked mutations. Journal of Neurochemistry 88, 401–410.

Li, X., Kumar, Y., Zempel, H., Mandelkow, E.M., Biernat, J., Mandelkow, E., 2011. Novel diffusion barrier for axonal retention of tau in neurons and its failure in neurodegeneration. EMBO Journal 30, 4825–4837.

Maday, S., Twelvetrees, A.E., Moughamian, A.J., Holzbaur, E.L.F., 2014. Axonal transport: Cargo-specific mechanisms of motility and regulation. Neuron 84, 292–309.

Miles, C.E., Keener, J.P., 2017. Bidirectionality from cargo thermal fluctuations in motor-mediated transport. Journal of Theoretical Biology 424, 37–48.

Miles, C.E., Lawley, S.D., Keener, J.P., 2018. Analysis of nonprocessive molecular motor transport using renewal reward theory. SIAM Journal on Applied Mathematics 78, 2511–2532.

Misgeld, T., Schwarz, T.L., 2017. Mitostasis in neurons: Maintaining mitochondria in an extended cellular architecture. Neuron 96, 651–666.

Nath, S., Meuvis, J., Hendrix, J., Carl, S.A., Engelborghs, Y., 2010. Early aggregation steps in alpha-synuclein as measured by FCS and FRET: Evidence for a contagious conformational change. Biophysical Journal 98, 1302–1311.

Newby, J., Bressloff, P.C., 2010. Random intermittent search and the tug-of-war model of motor-driven transport. Journal of Statistical Mechanics-Theory and Experiment P04014.

Puthanveettil, S.V., Monje, F.J., Miniaci, M.C., Choi, Y., Karl, K.A., Khandros, E., Gawinowicz, M.A., Sheetz, M.P., Kandel, E.R., 2008. A new component in synaptic plasticity: Upregulation of kinesin in the neurons of the gill-withdrawal reflex. Cell 135, 960–973.

Raichur, A., Vali, S., Gorin, F., 2006. Dynamic modeling of alpha-synuclein aggregation for the sporadic and genetic forms of Parkinson’s disease. Neuroscience 142, 859–870.

Rosenberg, T., Gal-Ben-Ari, S., Dieterich, D.C., Kreutz, M.R., Ziv, N.E., Gundelfinger, E.D., Rosenblum, K., 2014. The roles of protein expression in synaptic plasticity and memory consolidation. Frontiers in Molecular Neuroscience 7, 86.

Roy, S., 2014. Seeing the unseen: The hidden world of slow axonal transport. Neuroscientist 20, 71–81.

Roy, S., 2016. Dynein’s life in the slow lane. Neuron 90, 907–909.

Roy, S., 2020. Finding order in slow axonal transport. Current Opinion in Neurobiology 63, 87–94.

Roy, S., Winton, M.J., Black, M.M., Trojanowski, J.Q., Lee, V.M.-., 2007. Rapid and intermittent cotransport of slow component-b proteins. Journal of Neuroscience 27, 3131–3138.

Roy, S., Winton, M.J., Black, M.M., Trojanowski, J.Q., Lee, V.M.Y., 2008. Cytoskeletal requirements in axonal transport of slow component-b. Journal of Neuroscience 28, 5248–5256.

Saha, A., Hill, J., Utton, M., Asuni, A., Ackerley, S., Grierson, A., Miller, C., Davies, A., Buchman, V., Anderton, B., Hanger, D., 2004. Parkinson’s disease alpha-synuclein mutations exhibit defective axonal transport in cultured neurons. Journal of Cell Science 117, 1017–1024.

Schapira, A. H. V., Lang, A. E. T., & Fahn, S. (Eds.). (2010). Movement disorders 4: Blue books of neurology series, volume 35. Philadelphia, PA: Saunders.

Scholz, T., Mandelkow, E., 2014. Transport and diffusion of tau protein in neurons. Cellular and Molecular Life Sciences 71, 3139–3150.

Scott, D.A., Das, U., Tang, Y., Roy, S., 2011. Mechanistic logic underlying the axonal transport of cytosolic proteins. Neuron 70, 441–454.

Shahmoradian, S.H., Lewis, A.J., Genoud, C., Hench, J., Moors, T.E., Navarro, P.P., Castano-Diez, D., Schweighauser, G., Graff-Meyer, A., Godie, K.N., Sutterlin, R., Huisman, E., Ingrassia, A., de Gier, Y., Rozemuller, A.J.M., Wang, J., De Paepe, A., Erny, J., Staempfli, A., Hoernschemeyer, J., Grosserueschkamp, F., Niedieker, D., El-Mashtoly, S.F., Quadri, M., Van IJcken, W.F.J., Bonifati, V., Gerwert, K., Bohrmann, B., Frank, S., Britschgi, M., Stahlberg, H., Van de Berg, W.D.J., Lauer, M.E., 2019. Lewy pathology in Parkinson’s disease consists of crowded organelles and lipid membranes. Nature Neuroscience 22, 1099–1109.

Tang, Y., Das, U., Scott, D.A., Roy, S., 2012. The slow axonal transport of alpha-synuclein-mechanistic commonalities amongst diverse cytosolic cargoes. Cytoskeleton 69, 506–513.

Toba, S., Jin, M., Yamada, M., Kumamoto, K., Matsumoto, S., Yasunaga, T., Fukunaga, Y., Miyazawa, A., Fujita, S., Itoh, K., Fushiki, S., Kojima, H., Wanibuchi, H., Arai, Y., Nagai, T., Hirotsune, S., 2017. Alpha-synuclein facilitates to form short unconventional microtubules that have a unique function in the axonal transport. Scientific Reports 7, 16386.

Twelvetrees, A.E., Pernigo, S., Sanger, A., Guedes-Dias, P., Schiavo, G., Steiner, R.A., Dodding, M.P., Holzbaur, E.L.F., 2016. The dynamic localization of cytoplasmic dynein in neurons is driven by kinesin-1. Neuron 90, 1000–1015.

Utton, M., Noble, W., Hill, J., Anderton, B., Hanger, D., 2005. Molecular motors implicated in the axonal transport of tau and alpha-synuclein. Journal of Cell Science 118, 4645–4654.

Wong, M.Y., Zhou, C., Shakiryanova, D., Lloyd, T.E., Deitcher, D.L., Levitan, E.S., 2012. Neuropeptide delivery to synapses by long-range vesicle circulation and sporadic capture. Cell 148, 1029–1038.

Yang, M., Hasadsri, L., Woods, W.S., George, J.M., 2010. Dynamic transport and localization of alpha-synuclein in primary hippocampal neurons. Molecular Neurodegeneration 5, 9.

Zadeh, K.S., 2008. Parameter estimation in flow through partially saturated porous materials. Journal of Computational Physics 227, 10243–10262.

Zadeh, K.S., 2011. A synergic simulation-optimization approach for analyzing biomolecular dynamics in living organisms. Computers in Biology and Medicine 41, 24–36.

Zadeh, K.S., Montas, H.J., 2014. Parametrization of flow processes in porous media by multiobjective inverse modeling. Journal of Computational Physics 259, 390–401.

Zadeh, K.S., Shah, S.B., 2010. Mathematical modeling and parameter estimation of axonal cargo transport. Journal of Computational Neuroscience 28, 495–507.

